# Modelling the distribution of *Aedes aegypti* and *Aedes albopictus* using climate, host density and interspecies competitive effects

**DOI:** 10.1101/498238

**Authors:** Bingyi Yang, Brooke A. Borgert, Barry W. Alto, Carl K. Boohene, Peter Brabant, Joe Brew, Kelly Deutsch, James T. DeValerio, Rhoel R. Dinglasan, Daniel Dixon, Joseph M. Faella, Sandra L. Fisher-Grainger, Gregory E. Glass, Reginald Hayes, David F. Hoel, Austin Horton, Agne Janusauskaite, Bill Kellner, Moritz U.G. Kraemer, Eric Leveen, Keira J. Lucas, Johana Medina, Rachel Morreale, William Petrie, Robert C. Reiner, Michael T. Riles, Henrik Salje, David L. Smith, John P. Smith, Amy Solis, Jason Stuck, Chalmers Vasquez, Katie F. Williams, Rui-De Xue, Derek A.T. Cummings

## Abstract

Florida faces the challenge of repeated introduction and autochthonous transmission of arboviruses transmitted by *Aedes aegypti* and *Aedes albopictus*. Empirically-based predictive models of the spatial distribution of these species would aid surveillance and vector control efforts. To predict the occurrence and abundance of these species, we fit mixed-effects zero-inflated negative binomial regression to a mosquito surveillance dataset with records from more than 200,000 trap days, covering 73% of the land area and ranging from 2004 to 2018 in Florida. We found an asymmetrical competitive interaction between adult populations of *Aedes aegypti* and *Aedes albopictus* for the sampled sites. Wind speed was negatively associated with the occurrence and abundance of both vectors. Our model predictions show high accuracy (72.9% to 94.5%) in the validation tests leaving out a random 10% subset of sites and data from 2018, suggesting a potential for predicting the distribution of the two *Aedes* vectors.

## INTRODUCTION

*Aedes* mosquitoes, in particular, *Aedes aegypti* (Linnaeus) and *Aedes albopictus* (Skuse), are the primary vectors of multiple arboviruses including dengue virus (DENV), Zika virus (ZIKV), yellow fever virus, and chikungunya virus (CHIKV)(Bargielowski and Lounibos, 2016; Kotsakiozi et al., 2017; Lounibos and Kramer, 2016). The incidence of these viruses in humans is driven, in part, by the close overlapping habitats of humans and these vectors (Charrel et al., 2014). In the absence of effective vaccines, reducing contact between mosquitoes and humans through targeted mosquito control is regarded as the most effective approach to reduce risk of mosquito-borne arbovirus transmission. There have been several efforts to create large-scale estimates of the spatial presence and abundance of these vectors using a variety of collection methods and data from literature reports and entomological surveys of mosquito occurrence. Global maps have been generated using climate and socio-economic variables, relying on a strong dependence of mosquito populations to temperature and rainfall (Brady et al., 2014; Kraemer et al., 2015b; Leta et al., 2018). These efforts have uncertainty associated with publication bias and variability of collection methods. Large-scale data collected by standardized surveillance methods could improve the certainty and precision of occurrence and abundance maps. Here, we use a dataset covering around 102,000 km# (73%) and more than 200,000 trap days spanning 17 years of observation (Figs 1 and S7). We built a mixed-effects zero-inflated negative binomial (ZINB) model to characterize and predict the occurrence and abundance of *Ae. aegypti* and *Ae. albopictus*, simultaneously using climate and human population density data.

**Figure 1.**
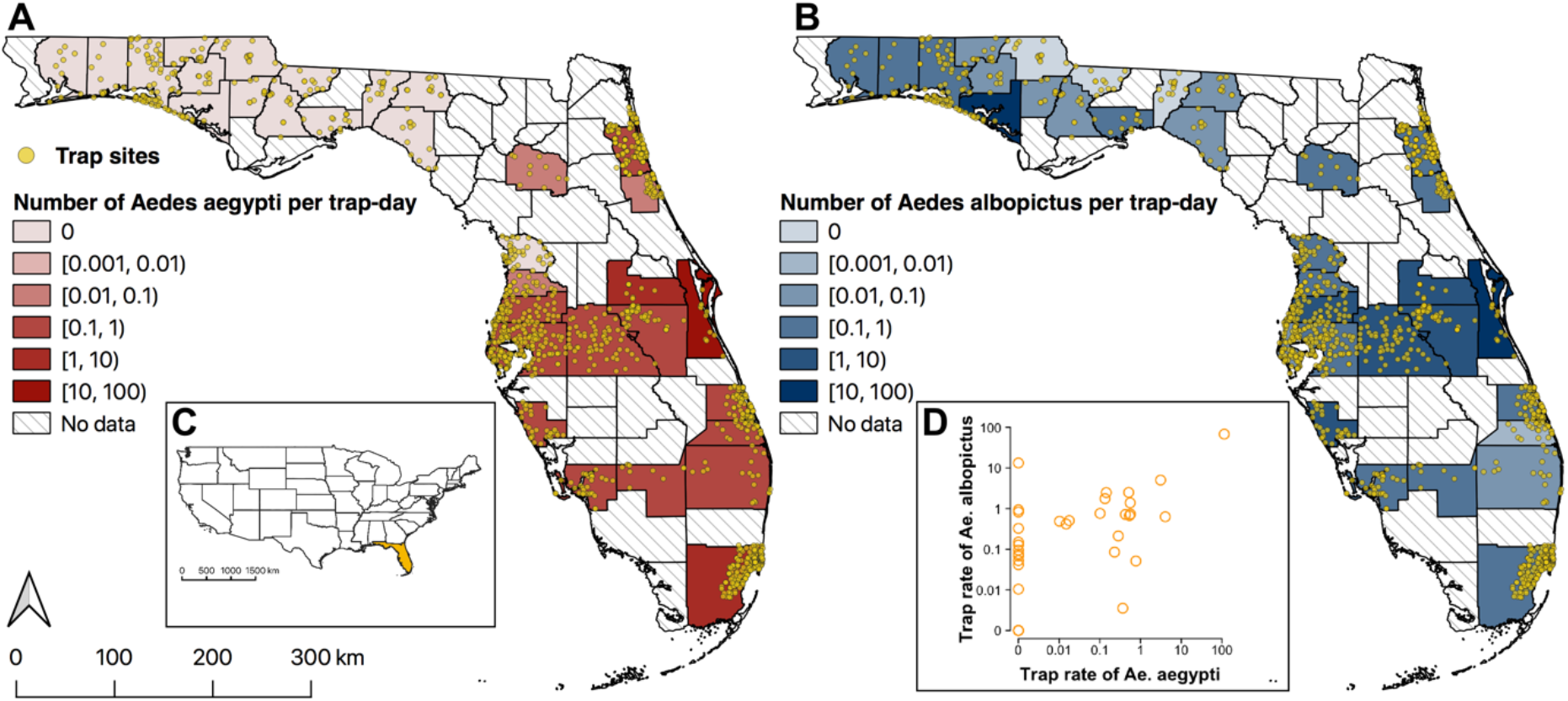
Locations of traps and geographic variation in abundance of *Aedes aegypti* (A) and *Aedes albopictus* (B) in Florida. Color (red for *Ae. aegypti*, blue for *Ae. albopictus*) indicates mean abundance per trap day in each county. Diagonal lines indicate counties without data. Inset (C) shows the location of Florida (orange) in the contiguous US. Plot (D) shows *Ae. aegypti* versus *Ae. albopictus* abundances in each county.

Florida has suffered from the introduction and autochthonous transmission of DENV (Muñoz-Jordán et al., 2013; Teets et al., 2014), CHIKV (Kendrick et al., 2014) and ZIKV (Grubaugh et al., 2017; Likos et al., 2016) and remain at high risk of transmission due to repeated pathogen introductions, high densities of *Ae. aegypti* and *Ae. albopictus* (Kraemer et al., 2015b) and favorable meteorological conditions (Grubaugh et al., 2017; Monaghan et al., 2016). Studies have shown a positive relationship between human Zika and dengue cases and larger *Ae. aegypti* populations in urban areas (Bowman et al., 2014; Grubaugh et al., 2017). Therefore, characterizing the population size of the two *Aedes* species over time and space could aid in examining the risk of local arbovirus transmission and spread in Florida and inform more effective and targeted mosquito control efforts.

Although coexistence of the two *Aedes* vectors is reported (Lounibos et al., 2016), declining populations and displaced habitats of *Ae. aegypti* have been observed in several places, including Florida (Bagny et al., 2009; Kaplan et al., 2010; Lounibos and Kramer, 2016; O’Meara et al., 1995). In particular, the habitats of *Ae. aegypti* were restricted to urban areas while those of *Ae. albopictus* were found to increase in suburban and rural areas in Florida (Lounibos, 2002). The proposed mechanisms for the displacements of *Ae. aegypti* include species interactions such as the superiority of *Ae. albopictus* to compete for resources at the larval stage and asymmetric sterilization at the adult stage after interspecific mating which favors *Ae. albopictus* (Bargielowski and Lounibos, 2016; Juliano, 2009; Lounibos and Kramer, 2016). Previous studies modelled the current spatial distribution of *Ae. aegypti* and *Ae. albopictus* by applying boosted regression trees to a comprehensive global database of *Aedes* occurrence (Kraemer et al., 2015a, 2015b) and characterized the spatial and temporal abundance of the two *Aedes* species in a local southern Florida county (Reiskind and Lounibos, 2013). However, these studies, which estimated the distribution and abundance of *Aedes*, are limited because of the minimal amount of data from standardized collections of mosquito populations and failure to consider the species interactions between *Ae. aegypti* and *Ae. albopictus* (Lounibos and Juliano, 2018). Additionally, inconsistent findings on the associations between their distribution and meteorological factors were reported according to a recent systematic review (Sallam et al., 2017).

The objective of this study was to simultaneously characterize the occurrence and abundance of the *Ae. aegypti* and *Ae. albopictus* mosquitoes using the routine mosquito surveillance data in Florida. To estimate if mosquitoes were present or not and if present, the number of adults in each trap location, a mixed-effects zero-inflated negative binomial (ZINB) regression was performed. Different predictors or factors were examined, like climate and human population density covariates, and the potential interaction between *Ae. aegypti* and *Ae. albopictus* based on their spatial and temporal abundance. Predictions on occurrence of *Ae. aegypti* and *Ae. albopictus* from models were assessed with and without abundance information to determine if real-time predictions based solely on climate data and human population density information provided accurate predictions.

## RESULTS

In total, the longitudinal training dataset included132,088 weekly records from 1,246 unique sites for *Ae. aegypti* and *Ae. albopictus*, respectively, covering 33 out of 67 counties. The dataset includes 53% of the land area in Florida and from 2004 to 2018 (Table 1, Figs. S1, S2 and S5). Traps were typically set for one day but a minority of collaborators reported counts from a trap that was set for multiple days (7.4%). Approximately 87.4% and 84.8% of trap episodes reported no adults collected for *Ae. aegypti* or *Ae. albopictus*, respectively. The majority (81.4%) of traps used were light traps, and the remaining 7.3% and 11.3% of traps used were BG Sentinel traps or other mosquito traps (Table S4), respectively (Table 1) (see more details for traps in the supplementary). A wider range and higher trap rate was reported for *Ae. albopictus* compared to *Ae. aegypti* in Florida, and as expected from previous studies, most *Ae. aegypti* were reported in southern Florida (Figs. 1 and S1). Both *Ae. aegypti* and *Ae. albopictus* were trapped more often between May to October (Fig. S1 and S2). The median human population density of the locations where the traps were set is 480.8 persons per km^2^ (Interquartile range (IQR), 112.5 to 1165.2 km^2^) (Fig S3). The median weekly average wind speed was 5.4 meter per second (IQR, 4.5 to 6.6 m/s), and the median relative humidity was 76.7% (IQR, 73.1 to 80.1 %) (Fig S3). The minimum temperature of the trap episodes ranged from 18.7 to 25.8 °C with median of 23.0 °C. The median difference of predicted maximum temperature on minimum temperature was 0 °C (IQR, −0.5 to 0.5 °C) (Fig S3).

**Table 1.**
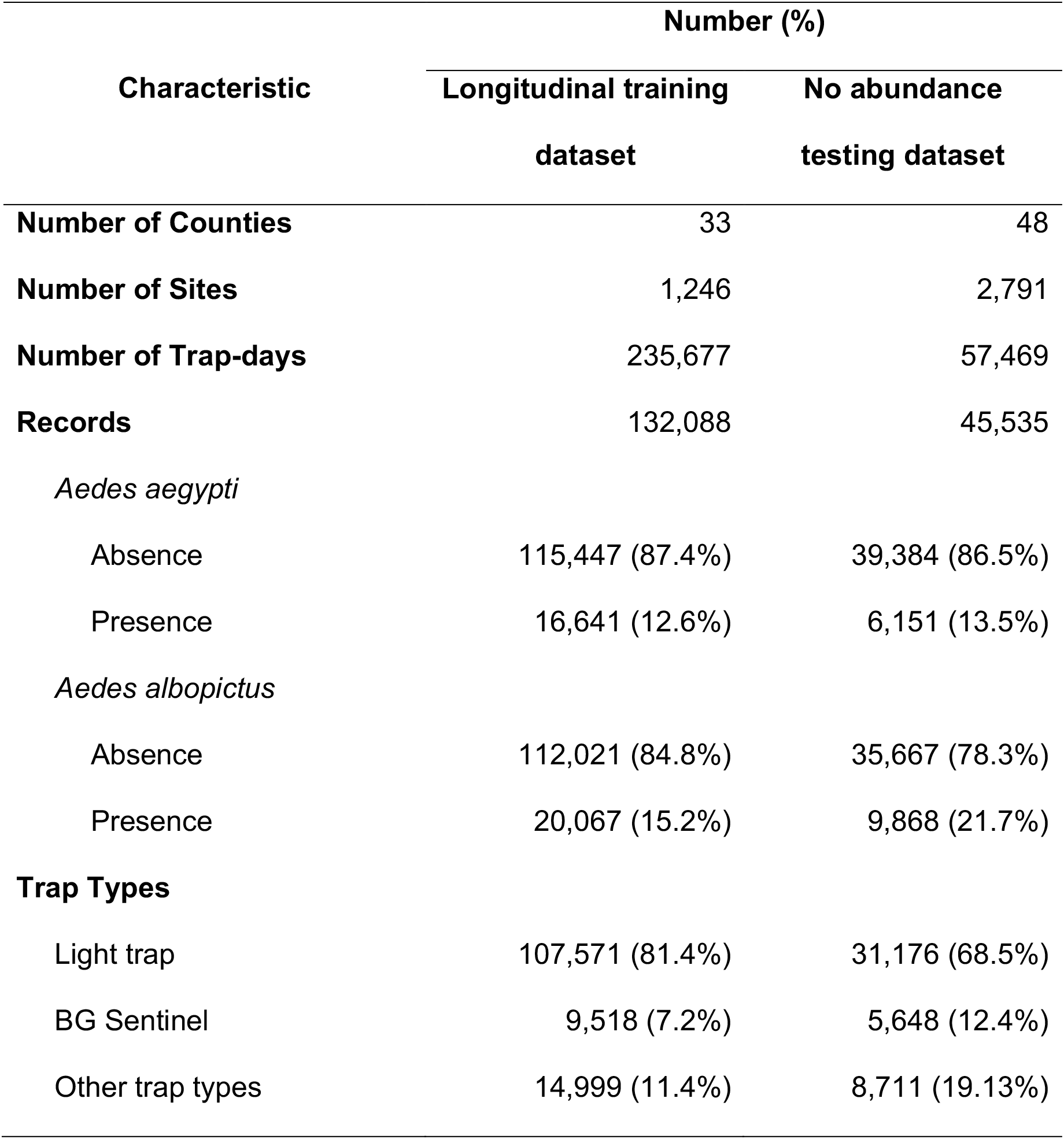
Characteristics of surveillance of *Aedes aegypti* and *Aedes albopictus* in Florida, 2004-2018.

### Presence and abundance of *Aedes*

The results from ZINB regression suggested the probability of presence of *Ae. aegypti* and *Ae. albopictus* in the current week was positively associated with the previous presence of its own species and the other species (Table 2). The abundance of both *Aedes* species was more likely to be higher if a higher abundance was reported for its own species (e.g. Incidence rate ratio (IRR) 1.03 and 1.02 for one week prior for the two vectors, respectively) (Table 2). The abundance of *Ae. aegypti* was negatively associated with the abundance of *Ae. albopictus* in the last three weeks (IRR: 0.992, 0.994 and 0.990 for one, two and three weeks earlier, respectively), while the abundance of *Ae. albopictus* seemed to be not associated with the previous abundance of *Ae. aegypti* (Table 2).

**Table 2.** Estimates of odds ratio (OR) and incidence rate ratio (IRR) from mixed-effects zero-inflated negative binomial analysis *Aedes aegypti* and *Aedes albopictus* in Florida, 2004-2018.

| Variables | *Aedes aegypti* |  | *Aedes albopictus* |  |
| --- | --- | --- | --- | --- |
|  | OR<br>(95% CrI<sup>†</sup>) | IRR<br>(95% CrI<sup>†</sup>) | OR<br>(95% CrI<sup>†</sup>) | IRR<br>(95% CrI<sup>†</sup>) |
| **Previous *Ae. aegypti* abundance/presence** |  |  |  |  |
| Trap rate in week *t*-1 | 2.46 (2.19, 2.76)* | 1.03 (1.02, 1.03)* | 1.21 (1.10, 1.34)* | 1.00 (1.00, 1.01) |
| Trap rate in week *t*-2 | 2.36 (2.11, 2.65)* | 1.03 (1.03, 1.03)* | 1.42 (1.28, 1.57)* | 1.00 (0.99, 1.00) |
| Trap rate in week *t*-3 | 1.84 (1.64, 2.07)* | 1.02 (1.01, 1.02)* | 1.04 (0.94, 1.15) | 1.00 (1.00, 1.01) |
| **Previous *Ae. albopictus* abundance/presence** |  |  |  |  |
| Trap rate in week *t*-1 | 1.30 (1.16, 1.47)* | 0.99 (0.99, 1.00)*† | 2.48 (2.32, 2.65)* | 1.02 (1.02, 1.03)* |
| Trap rate in week *t*-2 | 1.44 (1.29, 1.62)* | 0.99 (0.99, 1.00)*† | 2.19 (2.05, 2.35)* | 1.02 (1.01, 1.02)* |
| Trap rate in week *t*-3 | 1.28 (1.14, 1.44)* | 0.99 (0.99, 1.00)*† | 1.68 (1.57, 1.80)* | 1.02 (1.01, 1.02)* |
| **Human population density (100 persons/km<sup>2</sup>)** | 1.05 (1.03, 1.07)* | 1.00 (0.99, 1.02) | 0.95 (0.94, 0.97)* | 0.98 (0.97, 1.00)*† |
| **Meteorology** |  |  |  |  |
| Average wind speed (m/s) | 0.98 (0.95, 1.01) | 0.97 (0.96, 0.99)* | 0.97 (0.95, 0.99)* | 0.97 (0.95, 0.98)* |
| Minimum temperature (°C) | 1.01 (0.99, 1.02) | 1.13 (1.12, 1.14)* | 1.08 (1.07, 1.09)* | 1.09 (1.08, 1.10)* |
| Maximum temperature (°C) | 1.12 (1.03, 1.21)* | 0.91 (0.87, 0.95)* | 1.01 (0.95, 1.06) | 1.04 (1.00, 1.08)*† |
| Relative humidity (%) | 1.01 (1.00, 1.02) | 0.99 (0.98, 1.00)*† | 0.99 (0.98, 0.99)* | 1.00 (0.99, 1.00) |
| **Trap type** |  |  |  |  |
| BG sentinel | Ref. | Ref. | Ref. | Ref. |
| Light trap | 0.00 (0.00, 0.01)* | 0.40 (0.31, 0.51)* | 0.77 (0.60, 1.00)*† | 0.29 (0.24, 0.36)* |
| Other | 0.01 (0.00, 0.02)* | 0.20 (0.14, 0.29)* | 1.77 (1.28, 2.44)* | 0.25 (0.19, 0.33)* |
| **Random effects** |  |  |  |  |
| Site | 1.34 | 1.67 | 1.40 | 0.90 |
| County | 12.27 | 2.82 | 6.59 | 1.56 |
| **Dispersion parameter** | -- | 1.46 (1.42, 1.51) | -- | 1.13 (1.10, 1.17) |

\* P < 0.05. † Credible interval. ‡ The values with three effective digits for these estimates are (from right to left by row): 0.992 (0.987, 0.998), 0.994 (0.988, 0.999), 0.990 (0.985, 0.996), 0.984 (0.969, 0.998), 1.041 (1.001, 1.083), 0.986 (0.979, 0.994) and 0.775 (0.600, 0.999).

We found both the presence (Odds ratio (OR): 0.98, 0.95 to 1.01 and 0.97, 0.95 to 0.99, respectively) and abundance (IRR: 0.97, 0.96 to 0.99 and 0.97, 0.95 to 0.98, respectively) of Ae. *aegypti* and *Ae. albopictus* were negatively associated with the average wind speed of the week. Minimum temperature was positively associated with the occurrence (OR: 1.01 for Ae. *aegypti* and 1.08 for *Ae. albopictus*) and the abundance (IRR: 1.13 and 1.09 respectively) of both species. Maximum temperature was found to be negatively associated with the occurrence of Ae. *aegypti* (OR: 0.91, 0.87 to 0.95) but positively associated with the occurrence of Ae. *albopictus* (OR: 1.4, 1.00 to 1.08) (Table 2). We found the relative humidity was negatively associated with the abundance of *Ae. aegypti* (IRR: 0.99, 0.98 to 1.00) and the occurrence of *Ae. albopictus* (IRR: 0.99, 0.98 to 0.99). Model estimates using NOAA climate data were similar with our main results, except for the positive associations between maximum temperature and the abundance and presence for both species (Table S1). Greater precipitation was positively associated with the abundance for both *Ae. aegypti* (IRR: 1.42, 1.26 to 1.59) and *Ae. albopictus* (IRR: 1.09, 0.99 to 1.20), but not associated with the probability of presence (OR: 0.85, 0.69 to 1.05 and 1.5, 0.94 to 1.19, respectively) (Table S1).

Both the probability of presence (OR: 0.95, 0.94 to 0.97) and abundance (IRR: 0.98, 0.97 to 1.00) of *Ae. albopictus* were negatively associated with a higher human population density, while the probability of the presence of *Ae. aegypti* was positively associated with human population density (OR: 1.05, 1.03 to 1.07). We also found substantial heterogeneities of presence and abundance of these two *Aedes* species across trap sites and counties (Table 2). The greatest heterogeneity was found with the presence at a county level for both *Ae. aegypti* (random effects (RE): 12.27) and *Ae. albopictus* (RE: 6.592).

### Performance of model fits to the longitudinal training dataset

We compared the predictions from the main ZINB model with observed presence and abundance from the longitudinal training datasets (Fig. 2 and Fig. S6). Overall, our model fits well with both the occurrence and abundance estimates for *Ae. aegypti* and *Ae. albopictus* (Fig. 2). We observed that 91.1% and 84.9% of the predicted presence was consistent with the observed presence of *Ae. aegypti* and *Ae. albopictus*, respectively. Similarly, 83.8% and 77.0% of the predicted abundance was correlated with the observations, while 90.1% and 86.5% of the predicted abundance differed by ±1 per trap day from the observations (Fig. 2C-D). The values of Moran’s I are 0.47 (p < 0.01) and 0.08 (p = 0.02) for *Ae. aegypti* and *Ae. albopictus*, respectively, and is −0.03 (p = 0.81) for *Ae. aegypti* after removing data from Miami-Dade. Temporal differences were relatively larger during May and September, when the observed average trap rates were also higher, for both species (Fig. S6).

**Figure 2.**
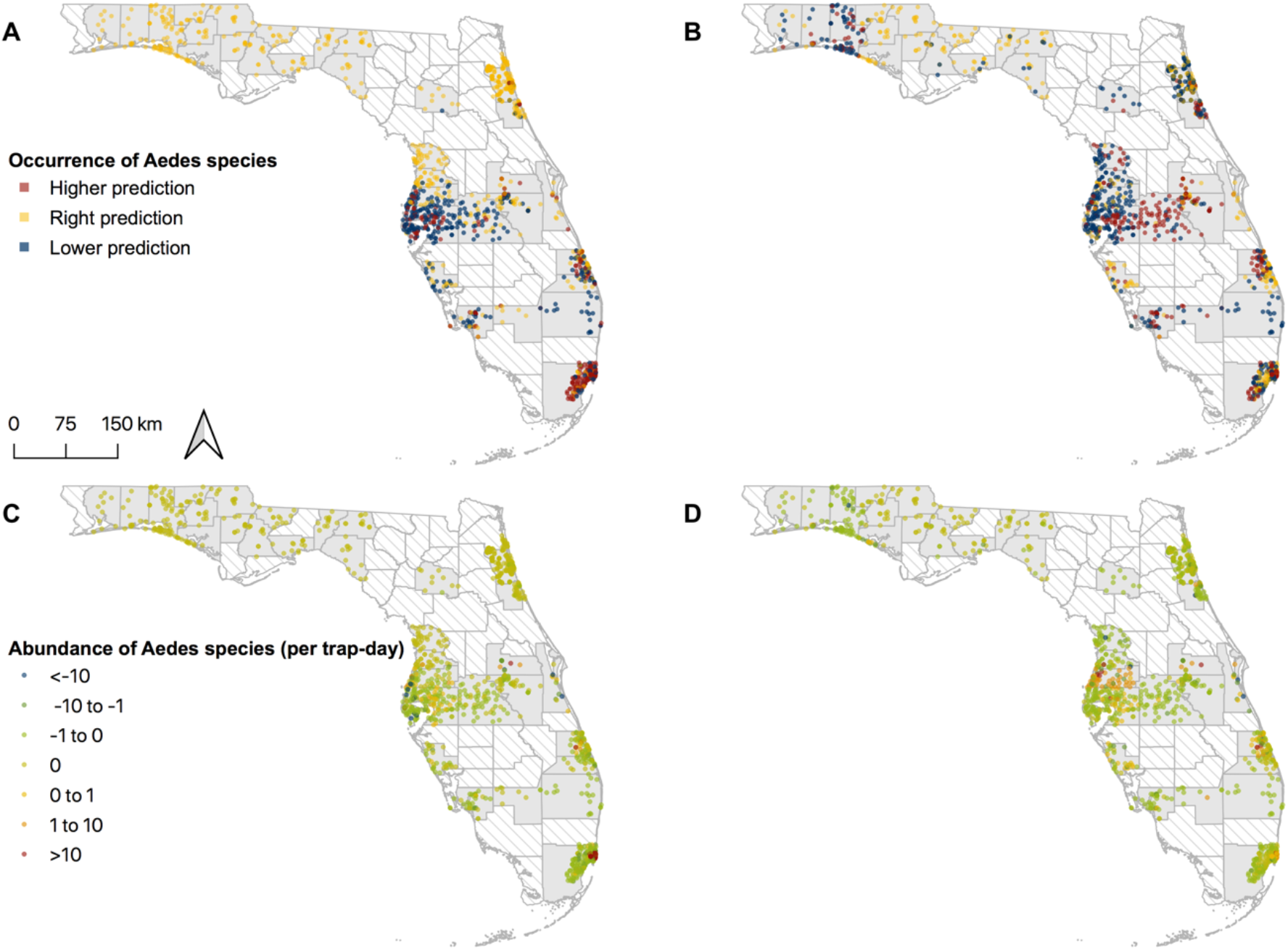
Geographic variation in model predictions in occurrence and abundance of *Aedes aegypti* and *Aedes albopictus*. (A) Occurrence of *Ae. aegypti*. (B) Occurrence of *Ae. albopictus*. (C) Abundance of *Ae. aegypti*. (D) Abundance of *Ae. albopictus*. Average difference between predictions and observations was calculated for each trap site.

### Performance of model in validation sets

For both spatial (Fig. 3A) and temporal (Fig. 43) test datasets, the model predictions are highly consistent with the observed presence of both *Ae. aegypti* (AUC: 0.93 and 0.92 for spatial and temporal predictions, respectively) and *Ae. albopictus* (AUC: 0.84 and 0.81 for spatial and temporal predictions, respectively). Overall, 86.2% and 82.1% of the predicted abundance were consistent with the observations for the spatial prediction of *Ae. aegypti* and *Ae. albopictus* (Fig. 3B-C), respectively, while 72.9% and 94.5% of the predictions were correct for the temporal predictions (Fig. 3E-F).

**Figure 3.**
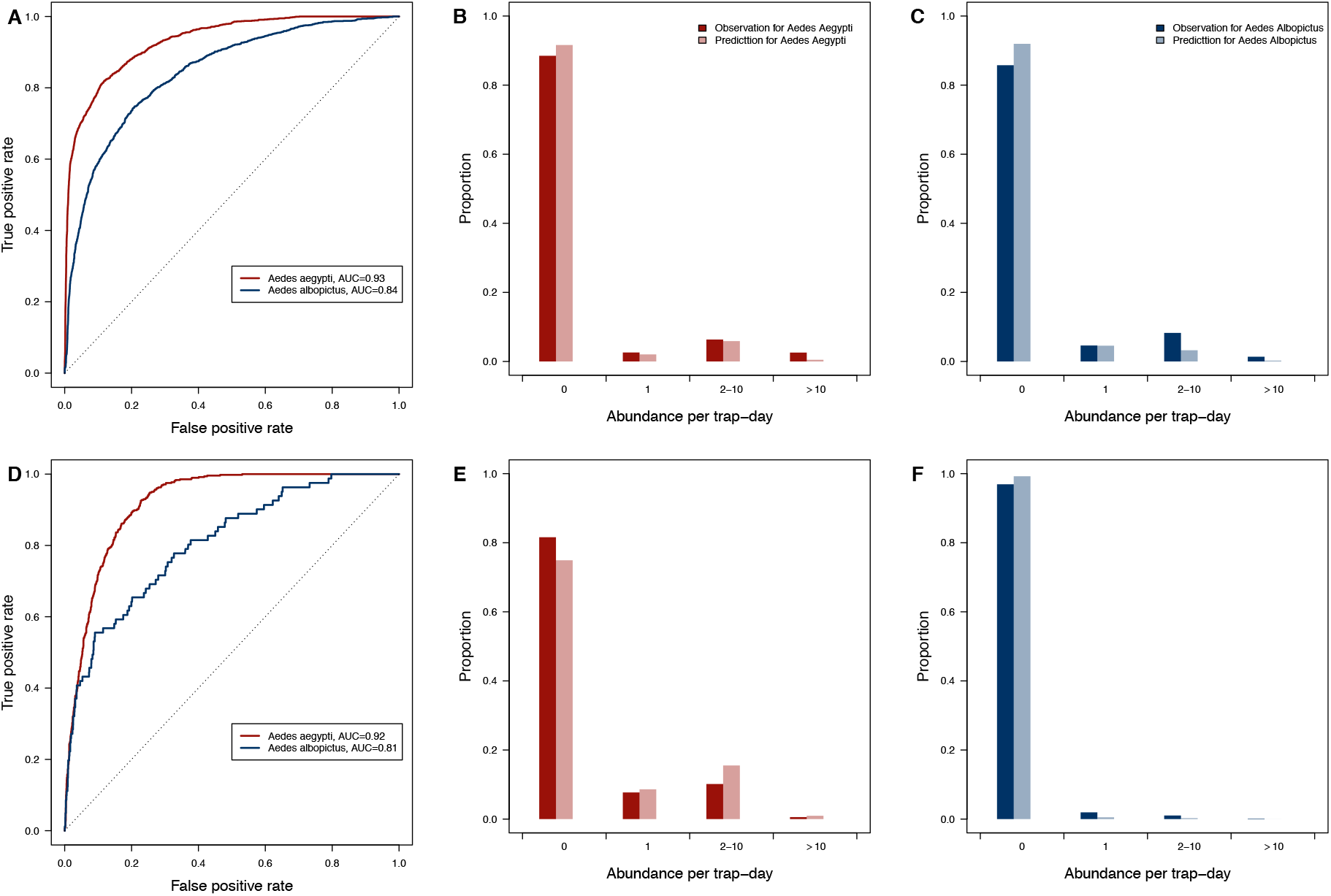
The performance of predictions in occurrence and abundance of *Aedes aegypti* and *Aedes albopictus*. (A-C) Records from 10% of trap sites were randomly selected as the test set and records from the rest traps were the train set. (D-F) Records from 2003 to 2016 were selected as the test set and records on and after 2017 were in the train set. The model was fit to the training set and predicted the test set.

We fit another ZINB model to the longitudinal training dataset without using information on the previous presence and abundance of both species, and applied the model to predict the no abundance testing dataset, which failed on the four consecutive four-week criteria. The no abundance testing dataset has total 45,535 trap episodes collected from 2,791 unique sites in 48 counties (Figs. S4 and S5).

The model provided good predictions in both presence (AUC: 0.90 for *Ae. aegypti* and 0.85 for *Ae. albopictus*) and abundance (82.8% and 70.2% right predictions, respectively) for the two *Aedes* species (Fig. S7).

Using our models, we predicted the number of *Ae. aegypti* and *Ae. albopictus* that would be expected to be found in traps in all points in the state. Fig. 4 shows predictions created using the “no abundance model” for August 1, 2018 incorporating random effects representing systematic differences in surveillance by county (Fig. 4 A and B) and only incorporating fixed effects (Fig. 4 C and D). BG traps were assumed to be used for all sites. Predictions in Fig. 4 present our estimates attempting to eliminate the impact of systematic differences in surveillance.

**Figure 4.**
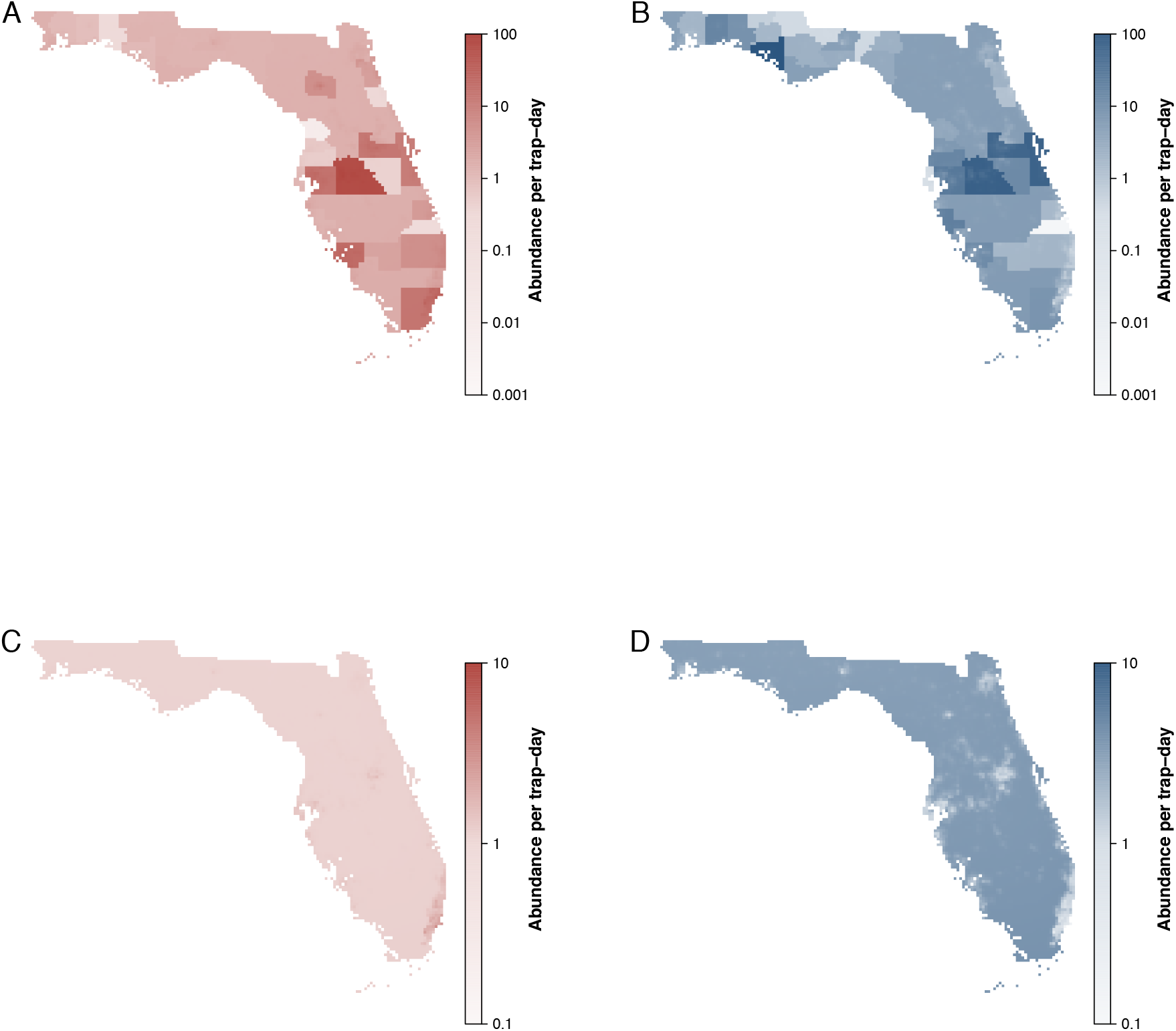
Maps of predicted counts of *Aedes aegypti* (red, A and C) and *Aedes albopictus* (blue, B and D) in August 1, 2018 in Florida. Predictions are derived from “no abundance model”. Parts A and B show results incorporating random effects representing differences in trapping counts by county. Parts C and D show results only incorporating fixed effects.

## DISCUSSION

We built models using more than 132,000 routine mosquito surveillance records from 33 counties in Florida collected from 2004 to 2018, to characterize and predict the occurrence and abundance of *Ae. aegypti* and *Ae. albopictus*. Our model performed well, particularly considering the stochastic nature of mosquito populations, trap efficiency and small-scale trap locations. We modelled random effects across sites and counties to account for inconsistencies and randomness and found the highest random effect was for the probability of presence at the county level, suggesting great heterogeneity of occurrence across counties possibly down to differences in surveillance and domestic mosquito control across counties.

Our results suggest a broad distribution of *Ae. albopictus* in Florida, while *Ae. aegypti* was more likely to be found in counties in southern Florida, a pattern similar to reports during the past two decades (Lounibos et al., 2016). This is also consistent with previous observations about the declining population of *Ae. aegypti* after the invasion of *Ae. albopictus* in the Southern United States (Bonizzoni et al., 2013; Lounibos, 2002). However, there is some evidence to suggest limited local recoveries of *Ae. aegypti* in relation to *Ae. albopictus*, in part, attributable to evolution of resistance to satyrization (Bargielowski et al., 2013; Bargielowski and Lounibos, 2016; Hopperstad and Reiskind, 2016; Lounibos et al., 2016). Our findings on the positive association between the probability of presence of adult mosquitoes of the two *Aedes* species suggest their niches have some overlap particularly in urban areas (Lounibos and Juliano, 2018). This is supported by the observed coexistence of *Ae. aegypti* and *Ae. albopictus* in Florida (Bonizzoni et al., 2013; Lounibos and Kramer, 2016; Reiskind and Lounibos, 2013) and the similar breeding behavior of the two species (Hashim et al., 2018). We also found evidence of competitive interactions between the two species. The abundance of *Ae. aegypti* was negatively associated with the previous abundance of *Ae. albopictus*, with the greatest effect size observed for the abundance of *Ae. albopictus* during the previous three-week period. A previous study revealed the breeding preference of *Ae. aegypti* in habitats without *Ae. albopictus* (Hashim et al., 2018). Our findings support the hypothesis that the two *Aedes* species can coexist but the abundance of adult *Ae. aegypti* are suppressed due to its failure to outcompete at the larval stage and/or the impact of interspecific mating (Bargielowski et al., 2013; Juliano, 2009; Lounibos and Kramer, 2016). Evolution of resistance to interspecific mating (i.e., satyrization-resistance) in *Ae. aegypti* populations is likely to promote coexsistence.(Bargielowski and Lounibos, 2016) Future control efforts targeting the *Aedes* species, especially *Ae. albopictus*, need to take into account the risk of resurgence of *Ae. aegypti*, which has been documented in Brazil (Kotsakiozi et al., 2017), and can be possible in Florida considering recent reports of the rapid evolution of satyrization-resistant *aegypti* (Bargielowski and Lounibos, 2016), coupled with an observed increased in insecticide resistance as compared to Ae. *albopictus* (Estep, et al., unpublished) (“Distribution Maps – Florida Mosquito Information,” n.d.).

We found the presence and abundance of *Ae. albopictus* are negatively associated with human population density, while the presence of *Ae. aegypti* was positively associated with the human population density, which matches with reports that anthropophilic *Ae. aegypti* are more likely to be found in urban areas and the *Ae. albopictus* has wider range of habitats including peri-urban, vegetated and rural areas (Lounibos and Kramer, 2016; Metzger et al., 2017), mostly due to its wide range of host preference and a greater adaptation to different climates (Bonizzoni et al., 2013). Land cover status, which is an important predictor of distribution of these species by other reports (Kraemer et al., 2015b; Rey et al., 2006), was not included in our main analysis as it may be associated with the human population density and the vast majority of the *Ae. aegypti* were collected in developed areas with a large human presence, consistent with other studies (Rodrigues et al., 2015; Tsai and Teng, 2016). The observed positive association between human and *Ae. aegypti* density has practical implications for targeted mosquito control because these areas represent the greatest risk for arboviral infections (e.g., dengue(Padmanabha et al., 2012)).

Major presence of both species between May to October has also been reported previously and corresponds to Florida’s rainy season and associated availability of breeding sites, and abiotic factors such as temperature (Reiskind and Lounibos, 2013). Related to this, the negative association between wind speed and the presence and abundance of both species analysis can include some explanations such as high wind speed hindering the effective trapping of the mosquitoes; therefore, traps are more likely to have no or fewer collections of mosquitoes during windy days. Also, mosquito activity and therefore host-seeking have been shown to be affected by higher wind speeds; presumably due to the combined effect of affected flight distance and pattern as well as the poor dispersal of the CO2 plume for both short and long distances.

Our results suggest positive associations between temperature and observed abundance of adult *Ae. aegypti* and *Ae. albopictus* when using the NOAA data, while inconsistent findings on the association between maximum temperature and the abundance of *Ae. aegypti* was found when using the NASA data. A study suggested higher tolerance of low temperature in adult *Ae. aegypti* compared to *Ae. albopictus*, leading to a relatively lower mortality of adult *Ae. aegypti* in low temperature and a milder effect of temperature on the presence of *Ae. aegypti* (Brady et al., 2013). One previous study observed that *Ae. albopictus* prefer to live in cooler areas in Florida (Bonizzoni et al., 2013). However, different local adaptations by these *Aedes* species to climatic changes were also reported both in and outside Florida (Lounibos and Kramer, 2016; Muttis et al., 2018). Despite these discussions with relation to habitat and mortality of the two *Aedes* vectors and temperature, seasonality can be used to predict the patterns of presence and abundance of these two *Aedes* species and the incidence of diseases transmitted by the these mosquito vectors (Monaghan et al., 2016; Reiskind and Lounibos, 2013; Xu et al., 2017).

We find a negative correlation between relative humidity the abundance of *Ae. aegypti* and the presence of *Ae. albopictus*. These findings support laboratory and field observations showing climate-driven egg mortality, with greater desiccation resistance of *Ae. aegypti* than *Ae. albopictus*, and species-specific responses in occupancy of containers with drier conditions favoring *Ae. aegypti* (Juliano et al., 2002; Lounibos et al., 2010; Mogi et al., 1996). Previous field studies have shown that dry periods are associated with disproportionately greater mortality *of Ae. albopictus* eggs than *Ae. aegypti* eggs in Florida (Juliano et al., 2002). Previous laboratory studies revealed desiccation stress on survival of adult *Ae. aegypti* and *Ae. albopictus* with mortality increasing non-linearly with decreasing relative humidity (Hylton, 1967; Lucio et al., 2013; Schmidt et al., 2018). The complex relation between adult survival and relative humidity and the observed disproportional distribution of higher relative humidity in Florida could drive the negative association (Fig S3). In addition, higher relative humidity was usually associated with greater precipitation, which was found to be positively correlated with the abundance of both vectors, but not the probability of occurrence of the two species in the sensitivity analysis (Table S2). The effect of precipitation on the abundance of these two *Aedes* species was considered to be mediated by induced egg hatching in containers upon flooding and promotion of vegetation after raining (Reiskind and Lounibos, 2013; Sallam et al., 2017). Larger effect of precipitation on the abundance of *Ae. aegypti* than of *Ae. albopictus* could be because the preference of the former to use breeding in artificial containers for development of the immature stages, which are prone to have more obvious influence from precipitation compared to vegetation.

The probability and efficacy of capturing *Ae. aegypti* and *Ae. albopictus* by a BG-sentinel trap was found to be greater compared to light traps (Table 2), which is consistent with previous findings (Li et al., 2016; Williams et al., 2006). We performed a sensitivity analysis by fitting a ZINB model to data collected by BG sentinel traps only and found the robustness of our main results are seemly unaffected not to be affected by the spatial distribution of BG sentinel traps (Fig. S8). In addition, we were not able to assess the role of attractants due to limited data available, which are believed to increase the capture efficacy of mosquitoes (de Ázara et al., 2013).

Our model which incorporates the previous abundance of heterospecific and conspecific *Aedes* species at a trap site demonstrates high accuracy in predicting the presence and abundance of *Ae. aegypti* and *Ae. albopictus* (Fig. 2 and Fig. S6). Analysis of long-term mosquito surveillance data is challenged by the excessive zero counts, which may be real absence, absence due to trap failure or adverse environmental conditions. The ZINB regression can model the two scenarios of absence simultaneously. However, variability in trap placement, efficiency of specific traps and other sources of variation in mosquito trapping practices may reduce our model performance. Performance tended to be lower when trapping rates were higher, while 97.1% (*Ae. aegypti*) and 96.8% (*Ae. albopictus*) of the differences between predicted and observed trap rate were within 5 per trap-day. A larger rate of inaccurate predictions was observed during months when trap rates of both mosquito species were higher, corresponding to the more dispersed variance of a higher trap rate. In addition, spatial autocorrelation was found for the model of *Ae. aegypti*, which was mainly due to the high autocorrelation between observations in Miami-Dade. The estimates and predictions are however not affected by the spatial autocorrelation, as suggested by the model fit to the longitudinal training dataset but removing data from Miami-Dade (Fig. S9 and Table S3).

Our model can be applied to predict spatial and temporal presence and abundance of *Ae. aegypti* and *Ae. albopictus* with good accuracy (Fig. 3). Although great variance was observed across counties and sites, we found that temporal prediction was more challenging for both species. Several sites first reported the occurrence or resurgence of *Ae. aegypti* in 2017, indicating the dynamic niches of the mosquitoes, which hinders the distribution forecasting of the two vectors.

We found that our model performed well in external validation even without recent data on each species (“no abundance model”). This suggests that accurate real-time forecasts could be generated without gathering and collating abundance data in realtime, giving timely predictions of the occurrence of *Ae. aegypti* and *Ae. albopictus* using only site-specific meteorological and human population density data. Results from the “no abundance model” incorporating fixed effects only provide homogenous predictions, which are largely informed by the human population density, while the empirical data however suggested great variations in the abundance captured across the counties (Fig. 1). This is could be partially due to the systematic differences in trapping practices and surveillance across counties and can be captured by the model incorporating random effects.

Many efforts have been made to map the distribution of *Ae. aegypti* and *Ae. albopictus* at broad regional scales, which were highly dependent on vegetation and meteorological factors (Brady et al., 2014; Kraemer et al., 2015b; Leta et al., 2018). Our study observes suppression of adult population of *Ae. aegypti* by *Ae. albopictus*, highlighting the importance of including species interactions in future mapping work as underscored by recent studies, especially when considering predictions at high spatial resolution (Lounibos and Juliano, 2018). Otherwise, the distribution of *Ae. aegypti* would likely be overestimated since the two *Aedes* vectors shared many common abiotic conditions. The median changes of predicted trap rate of *Ae. aegypti* in Miami-Dade are −17.0% (IQR: −21.0 to 19.3%) and −24.6% (IQR: −28.2 to 8.3%) when the trap rate for *Ae. albopictus* was 1 and 100 per trap-night, respectively. In addition, predictions from standardized longitudinal mosquito surveillance could aid to refine the distribution maps of these vectors by incorporating the seasonal pattern and real-time invasion activity. Finally, our empirical surveillance data can be used to validate and refine the local performance of large-scale maps. Integrating longitudinal surveillance could provide valuable information on absence and abundance, therefore reducing the sampling bias and disproportional weighting caused by presence only data (Wisz and Guisan, 2009).

There are several limitations to our study. First, our data has relatively more trap episodes during April to November, when the trap rate for these two vectors was often high. The estimated impact of low temperature on the presence and abundance of these two *Aedes* vectors may therefore be affected. Second, more than half of the records included in the main analysis are from Miami-Dade, St. Johns, Polk and Pinellas counties (Table S5). We have modelled the random effects across both sites and counties to account for the potential spatial variations of surveillance, which may improve the generalization capability of our conclusions. We were not able to characterize specific details of trap locations or other aspects of sites such as details of the built environment. These details might further improve forecasts.

Our models demonstrate potential for predicting the occurrence of *Ae. aegypti* and *Ae. albopictus*, to better inform targeted mosquito control efforts. Model predictions produced with and without the benefit of recent surveillance data were of high accuracy suggesting that real-time forecasts could be produced with just climate data alone.

## MATERIALS AND METHODS

### Mosquito surveillance data

Statewide surveillance data on 16 *Aedes* species was obtained by networking with Florida’s mosquito control districts, Clarke Scientific, the Florida Department of Agriculture Consumer Services, and the Florida Department of Health. Each control district is required to trap mosquitoes prior to conducting their control efforts by Florida Statutes 388 and 482. The traps were placed to acquire a representative sampling of the district including baseline traps placed in the same location annually, at risk areas due to environmental factors like increased standing water, locations within areas of known arbovirus transmission, and frequent areas of complaint. Information collected from these traps includes the speciated count and life phase of the trapped mosquitoes, date and duration of collection, type of trap, and coordinates of the trap sites. The collected mosquitoes were speciated according to standardized mosquito keys (Darsie and Morris, 2003). For missing data, the duration of collection was assumed to be one day, according to the common trapping practices, and coordinates were extracted from Google Maps based on the address of the site. The full dataset was aggregated to include data on adult *Ae. aegypti* and *Ae. albopictus*, two vectors related with arboviruses, on a week basis. The longitudinal training dataset for the zero-inflated negative binomial (ZINB) regression was extracted from the full dataset and included only data collected from sites with at least four consecutive weeks of surveillance and no missing explanatory variables.

### Abiotic variables

To examine the potential effects of meteorological factors on the trap rate of the two *Aedes* species, temperature (°C), wind speed (meter per second) and relative humidity (%) were included in the model. We obtained the daily meteorological data for Florida from the NASA Prediction of Worldwide Energy Resources(“NASA Prediction of Worldwide Energy Resources.,” n.d.) and applied the inverse distance weighting method (Pebesma and Gräler, 2014) to interpolate the daily weather raster of Florida with a 5 km ×5 km resolution. We also conducted a sensitivity analysis by using meteorological data from National Oceanic and Atmospheric Administration (NOAA).(National Oceanic and Atmospheric Administration, 2016) The weekly average of weather conditions was calculated as the mean of the weather conditions on the days the traps were collected. To account for the collinearity of the maximum and minimum temperature, we used the residuals of the linear regression of maximum temperature on minimum temperature as a proxy of the maximum temperature in the model, which was calculated as *ΔT_max_ = T_max_* − (*α* + *βT_min_*), where *T_max_* and *T_min_* denoting the observed maximum and minimum temperature, respectively, while *α* and *β* were estimated from the linear regression. The urbanization was modelled by including data on human population density, which was obtained from Center for International Earth Science Information Network with a 5 km ×5 km resolution (SEDAC and CIESIN, 2015). If the value was missing for a site, we extracted the corresponding environmental variables based on its coordinate and used the average drawn from a 5km buffer around the site.

### Statistical methods

We applied a ZINB regression model to the weekly abundance of *Ae. aegypti* and *Ae. albopictus* from the longitudinal training dataset, respectively, to account for the excessive zeros in the abundance data and the over-dispersed count of trapped mosquitoes, simultaneously. The ZINB model comprises a binary component (corresponding to the absence/presence of mosquitoes), and a negative binomial competent (corresponding to the abundance of mosquitoes). The estimates from the binary component (presented as odds ratio, OR) and the negative binomial component (presented as incidence rate ratio, IRR) represent the associations between the potential factors and the occurrence and abundance of these *Aedes* vectors, respectively. The potential factors included in the ZINB model for both species are: the previous abundance of *Ae. aegypti* and *Ae. albopictus* up to three weeks prior, weekly site-specific meteorological factors (i.e. wind speed, maximum and minimum temperature and relative humidity), human population density and type of mosquito traps. We examined the potential interaction between *Ae. aegypti* and *Ae. albopictus* by examining the relationship between the current abundance of one species with the previous abundance of another species. We used counts of each species detected in recent weeks to predict future weeks. To do this, we only considered records when data was available for four consecutive weeks prior. Trap type was included as an explanatory covariate as each of the traps used has a different effectiveness in trapping each species. We also included the random effects at both site level and county level, which were modelled for both components of ZINB model simultaneously. The detailed equations used for ZINB model are provided in the Supplementary information. Parameters were estimated by maximizing the likelihood using “glmmTMB” package (Brooks et al., 2017) in R version 3.5.0 (R Foundation for Statistical Computing, Vienna, Austria).

The model fitting was tested by comparing the observations with the predictions of occurrence and abundance from the longitudinal dataset. We assessed the spatial pattern by calculating the site-specific mean of residuals. The absence and presence were assigned as 0 and 1 respectively for calculation purposes. Moran’s I was calculated to assess the spatial autocorrelation (Bivand and Piras, 2015). We examined the temporal pattern of the model fitting by assessing the monthly 2.5% and 97.5% quantiles of the difference between the predicted and observed abundances for the two *Aedes species*.

We tested the prediction performance of the model both spatially and temporally. Prediction of a test set was based on a model fit from a training set and comparing the predicted and observed occurrence and abundance. In the spatial prediction, we randomly selected records from 127 (around 10% of total) sites to be the spatial testing set and used the records from the remainder of the sites as a spatial validation training set (Fig. S5). In the temporal prediction, we used data up to the year of 2017 as the temporal validation training set to predict data after 2017 (Fig. S5). The area under the receiver operating characteristic (AUC) was used to measure the performance of prediction on the mosquito occurrence. In addition, we also fit a ZINB model to the longitudinal training dataset without using information on the previous presence and abundance of *Ae. aegypti* and *Ae. albopictus* (“no abundance model”) and applied the “no abundance model” to an external no abundance testing dataset comprised of the surveillance records in the full dataset and failed on the four consecutive four-week criteria (Fig. S5).

## ACKNOWLEDGEMENTS

We thank Nathan Burkett-Cadena and Diana Araya Rojas for helpful discussion. We thank Adriane Rogers, Caitlin Gill, Caroline Efstathion, Frieda Lamberg, Ashley Pierre-Saint, Anthony Dennis, Taylor Thrail, Anthony Dennis, Cindy Mulla, Beth Kovach, Edfred Lontz, Pam Davis, Marah Clark, Matthew Mello, Bill Hockla, Toni Wright, Leonard Burns, Sherry Burroughs, Jaime Willoughby, Kylie E. Zirbel, Mitch Smeykal, Katie L Condra, Christopher Reisinger, Brian Lawton, Larry Gast, Kathleen Story, Wade Brenan, Richard Weaver, Gene Lamire, Terrill Mincey, James McNelly, Brenda Hunt and Amanda Baker for collecting and providing data. We thank Adam Shir, Yves Vaughan and Greg King for their help with digitizing the raw data. This project was supported by CDC Southeastern Center of Excellence in Vector-borne Diseases (CDC Cooperative Agreement U01CK000510). M.U.G.K. is supported by The Branco Weiss Fellowship - Society in Science, administered by the ETH Zurich and acknowledges funding from a Training Grant from the National Institute of Child Health and Human Development (T32HD040128) and the National Library of Medicine of the National Institutes of Health (R01LM010812, R01LM011965). Climate data were obtained from the NASA Langley Research Center POWER Project funded through the NASA Earth Science Directorate Applied Science Program.

The opinions expressed in the paper are those of the authors and do not necessarily represent the views of the United States Government.

## AUTHOR CONTRIBUTIONS

B.Y., B.A.B. and D.A.T.C. designed the study. B.Y., B.A.B. and D.A.T.C. wrote the first draft of the manuscript. B.W.A., R.R.D., G.E.G., M.U.G.K., R.C.R.Jr., H.S., and D.L.S. critically reviewed the manuscript. B.Y. performed the statistical analysis. B.Y. and B.A.B. collated the mosquito data. C.K.B., P.B., J.B., K.D., J.T.D., D.D., J.M.F, S.L.F., R.H., D.F.H., A.H., A.J., B.K., E.L., K.J.L., J.M., R.M., W.P., M.T.R., J.P.S., A.S., J.S., C.V., K.F.W. and R.D.X. provided and facilitated the collection of mosquito surveillance data. All authors contributed substantively to the revising and editing of the final draft.

## Conflict of interests

The authors declare no competing financial interests.

---

**Supplementary Figure S1.**
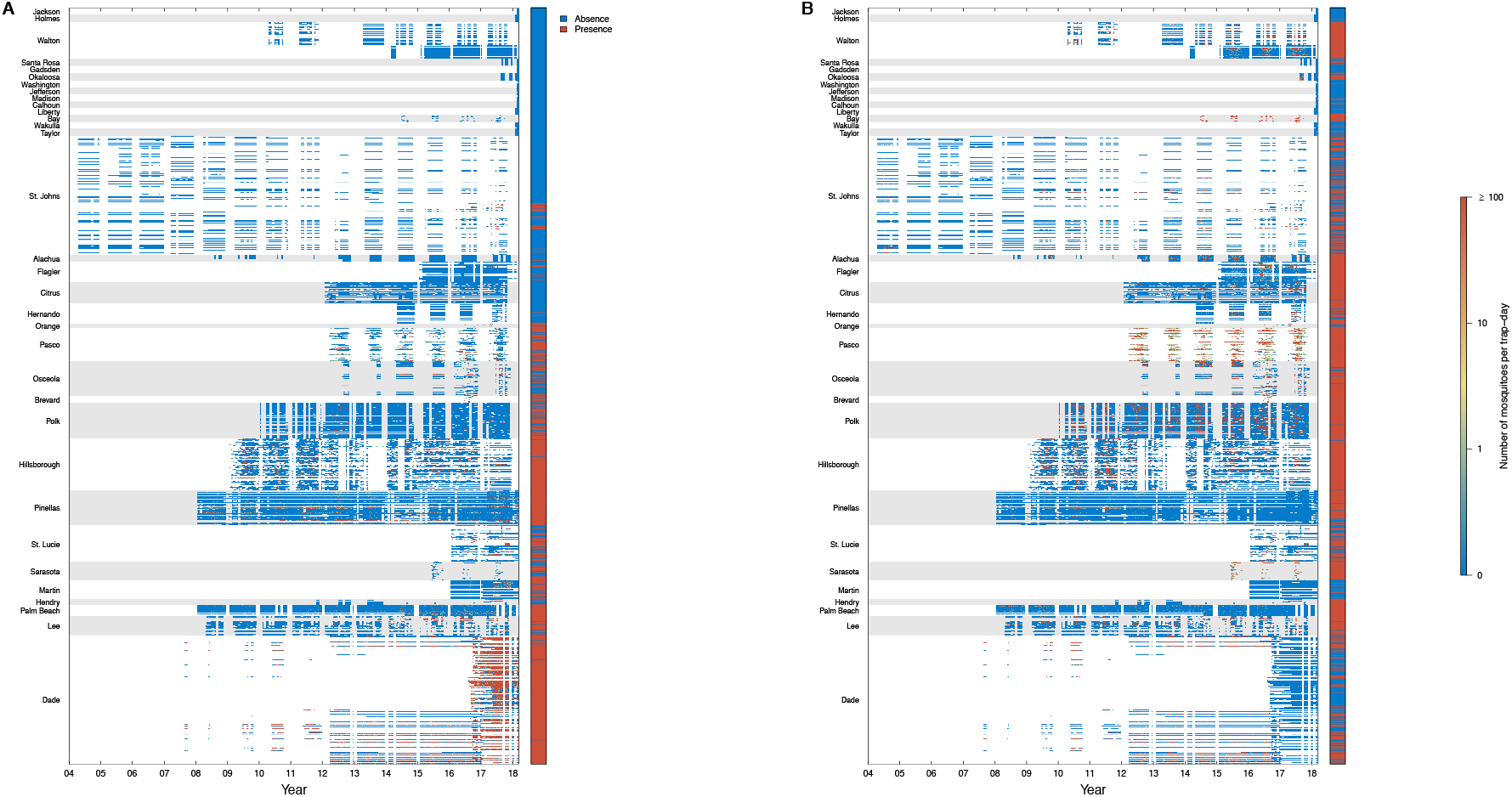
Spatial and temporal distribution of mosquito surveillance records. A, *Aedes aegypti*. B, *Aedes albopictus*. Trap sites and counties were ordered from north (upper) to south (lower). Heatmaps show weekly trap rate of each trap site. The sidebars indicate whether *Aedes aegypti* or *Aedes albopictus* had ever been reported by each site.

**Supplementary Figure S2.**
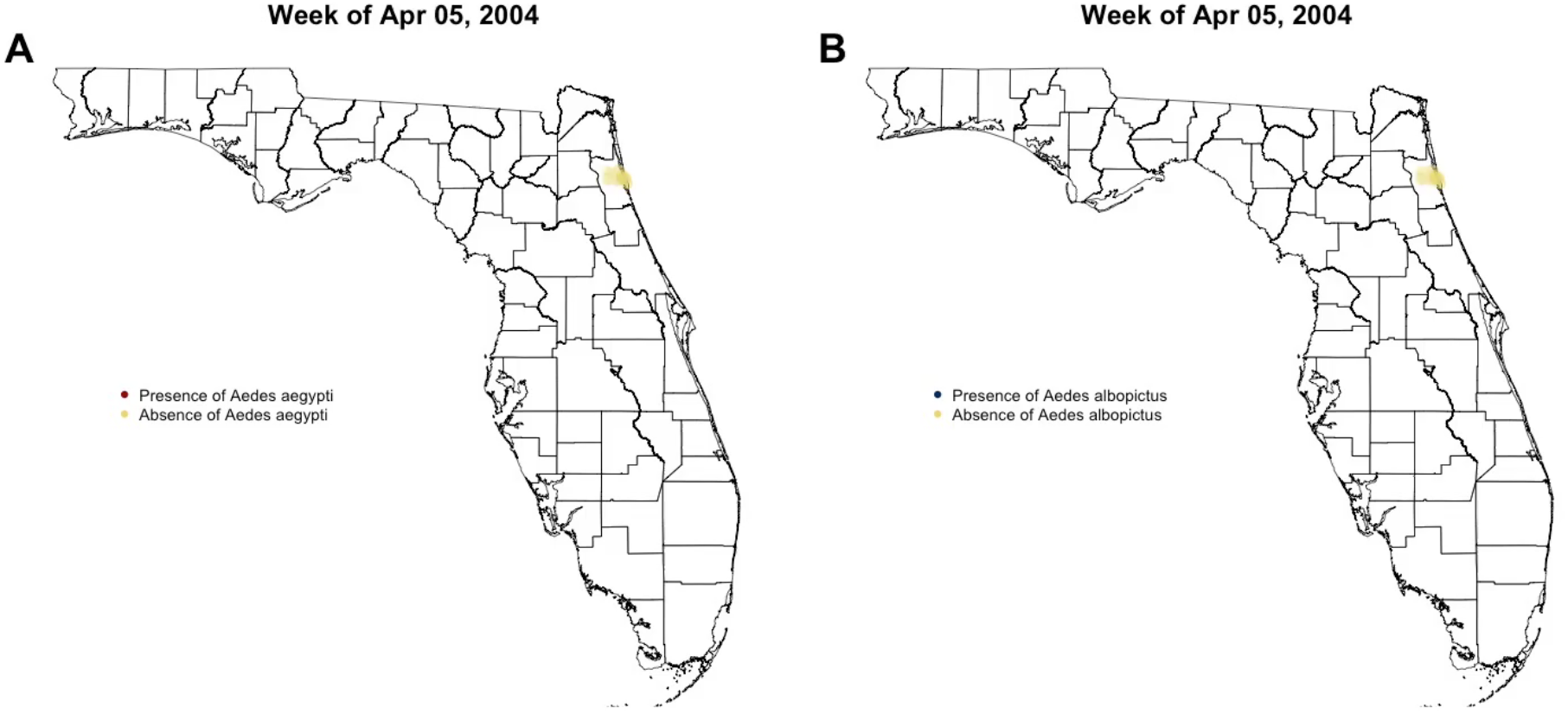
Weekly presence and absence of *Aedes aegypti* and *Aedes albopictus* in Florida.

**Supplementary Figure S3.**
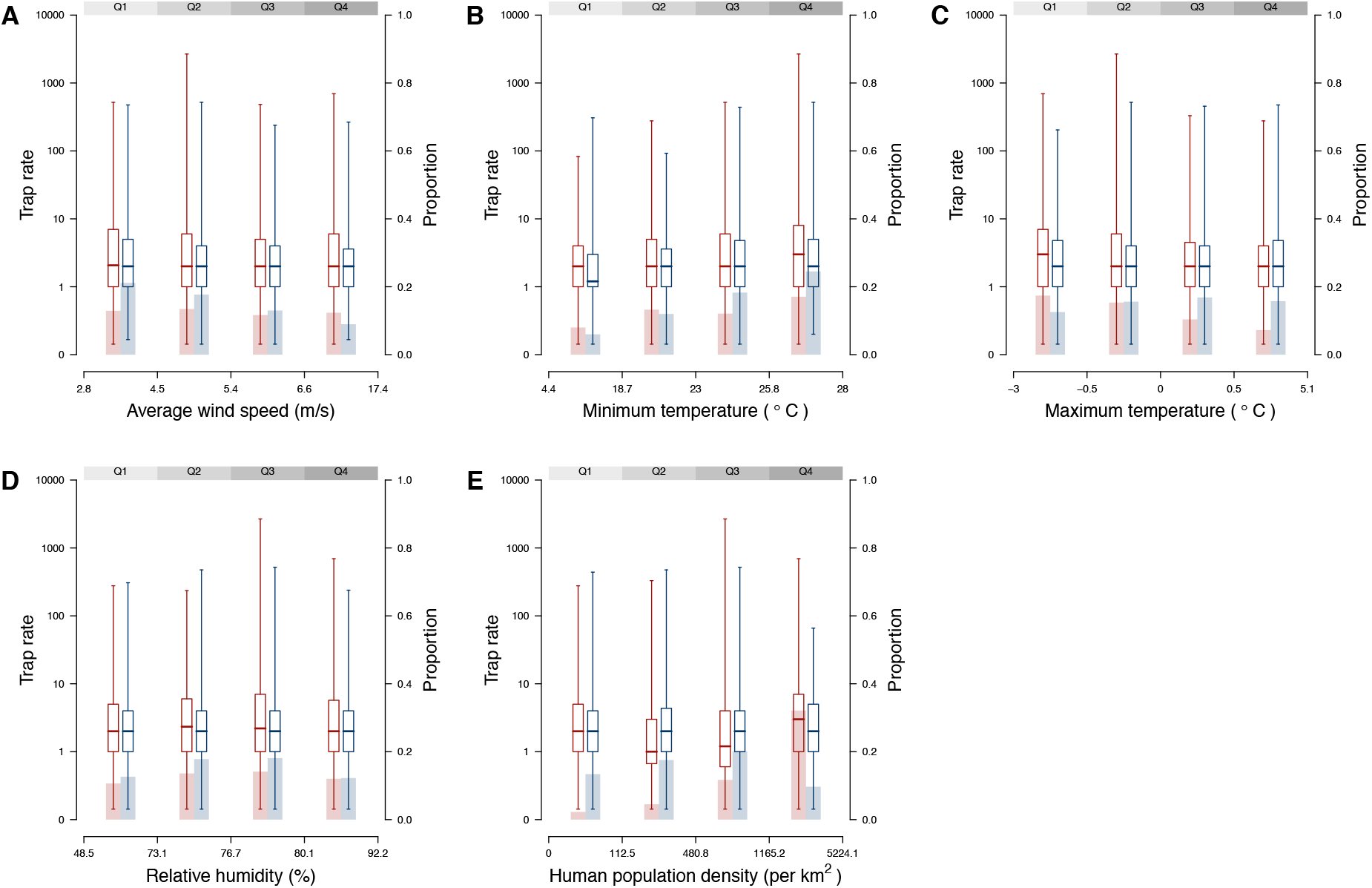
Relations between occurrence and abundance of *Aedes aegypti* and *Aedes albopictus* with abiotic variables. Values at x axis are the minimum, 25^th^ quantile, median, 75^th^ quantile and maximum value of the variable. Colored bar charts represent the proportion of occurrence reported by trap episodes. Colored box plots represent the median and interquartile range of the trap rate amongst traps where the vector occurred.

**Supplementary Figure S4.**
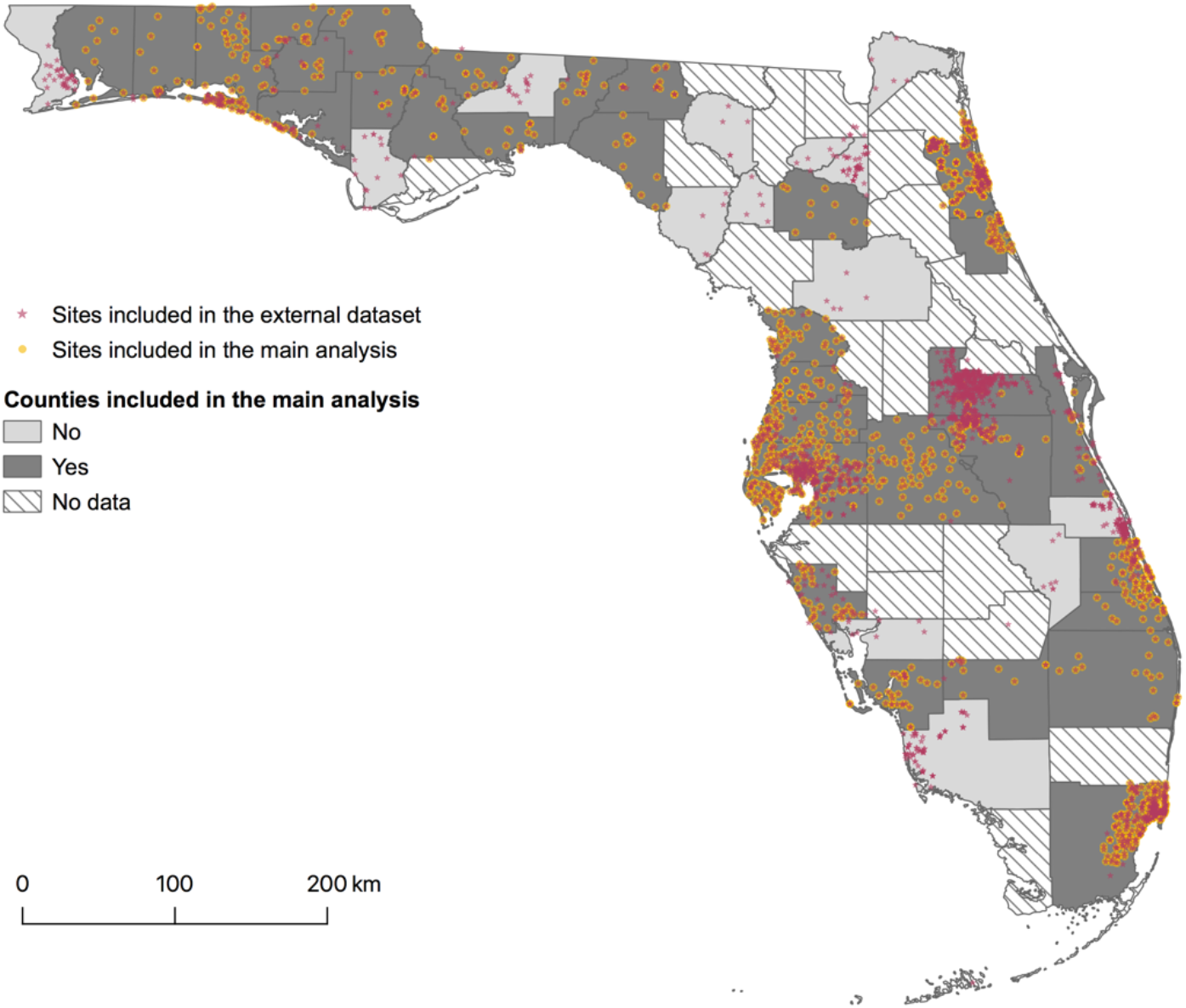
Comparison of trap locations by longitudinal training dataset and external no abundance testing dataset.

**Supplementary Figure S5.**
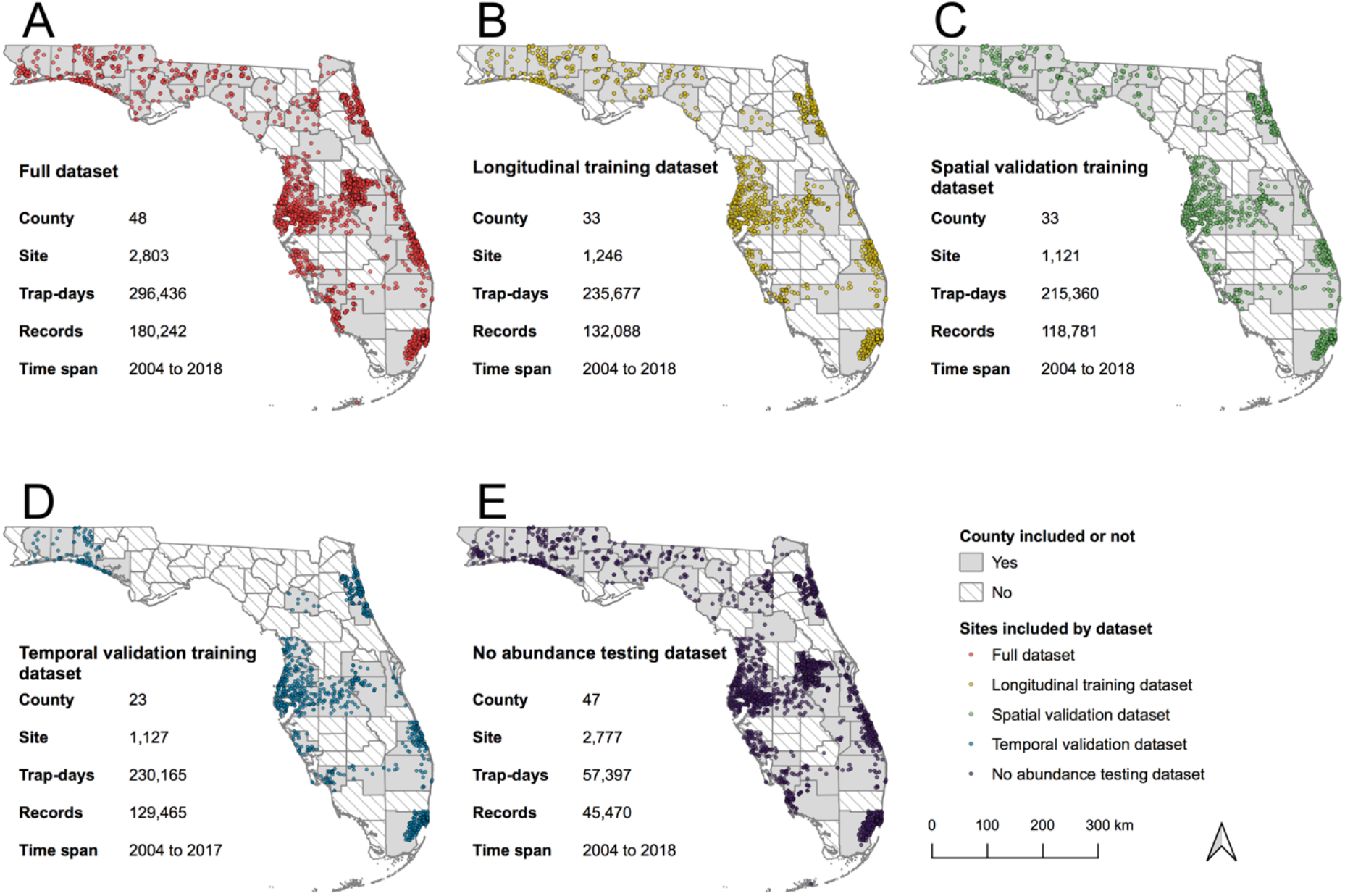
Comparison of five datasets used in the study.

**Supplementary Figure S6.**
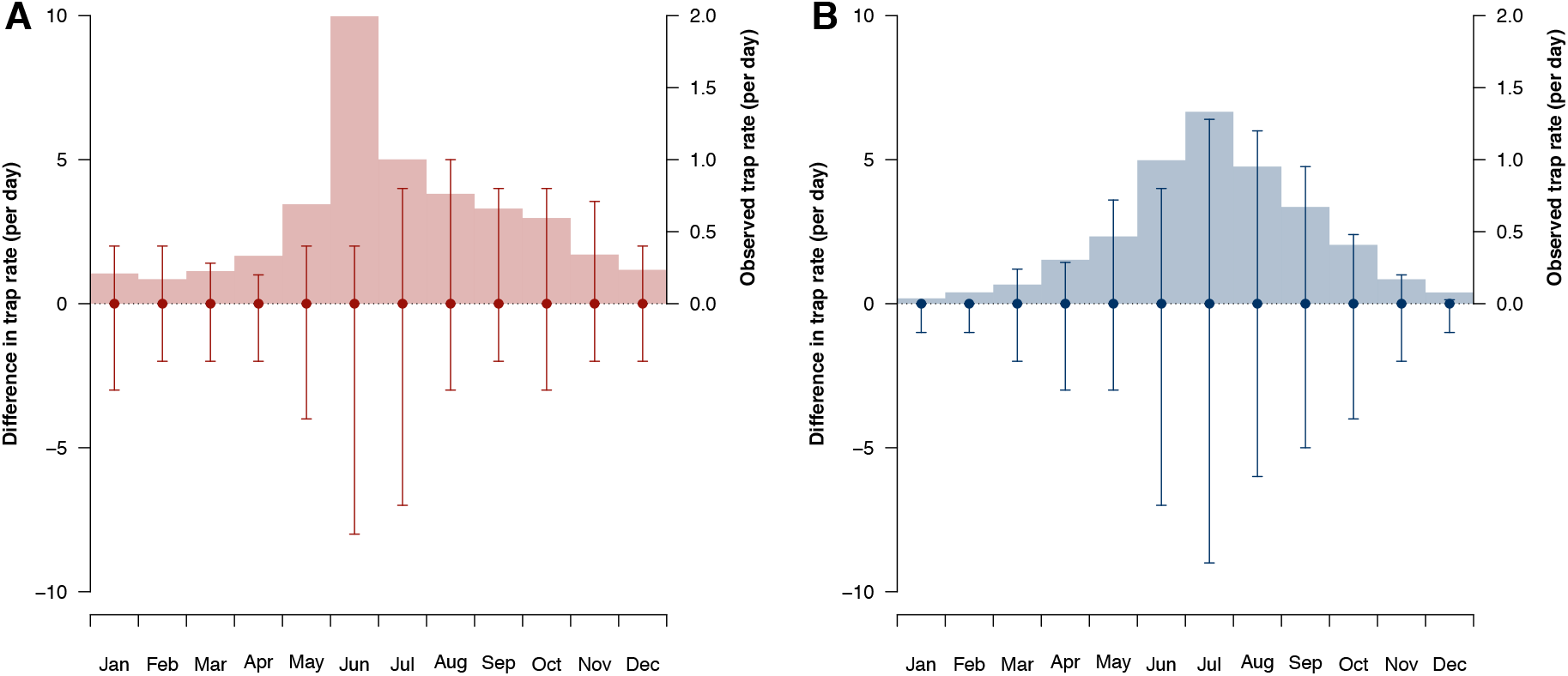
Temporal variation in model predictions in abundance of *Aedes aegypti* (A) and *Aedes albopictus* (B). Points are the median difference between predicted and observed abundance of *Aedes aegypti* and *Ae. albopictus* from the main analysis. Intervals are the 2.5% and 97.5% quantile of difference between predicted and observed abundance of the two *Aedes* species. Histograms are the monthly average of observed trap rates.

**Supplementary Figure S7.**
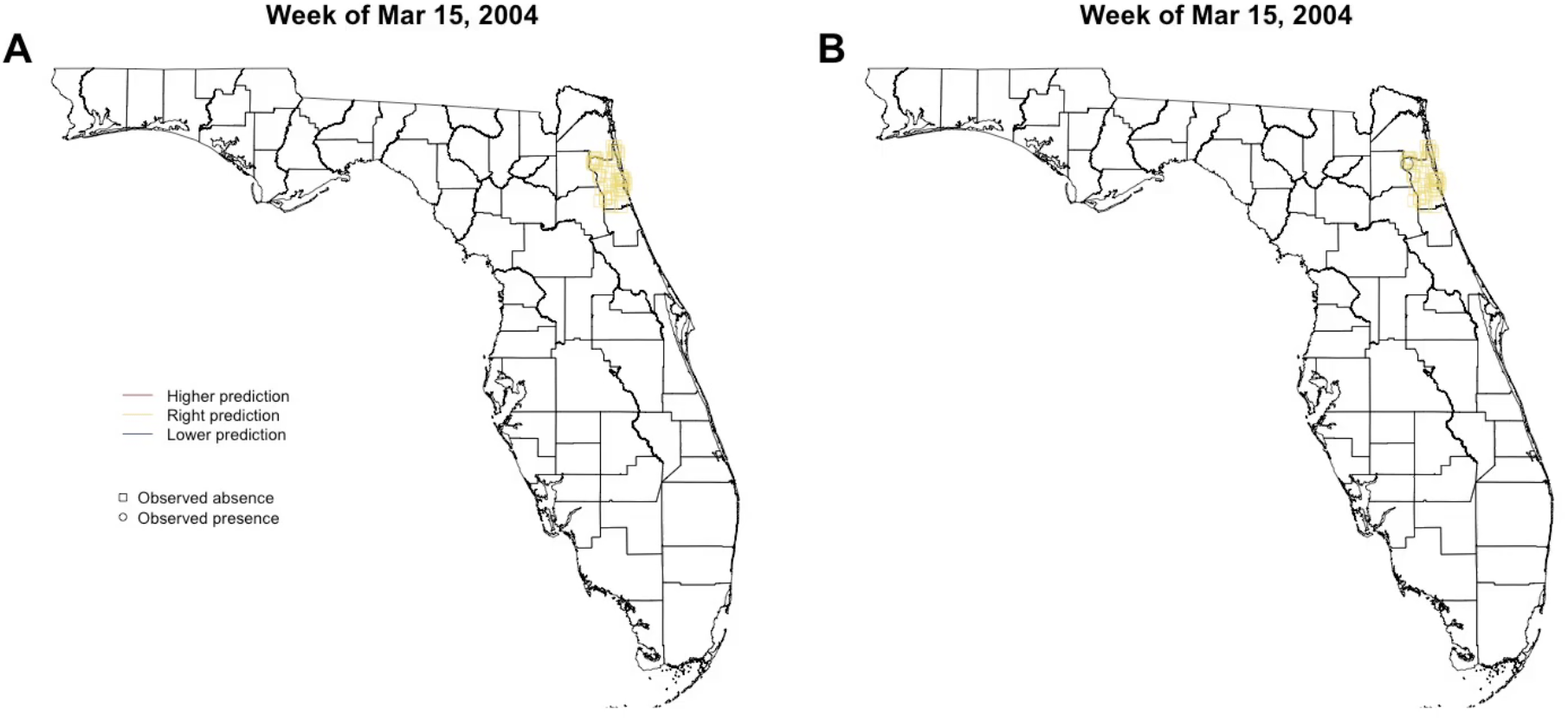
Weekly predictions of occurrence of *Aedes aegypti* and *Aedes albopictus* from no abundance model in Florida.

**Supplementary Figure S8.**
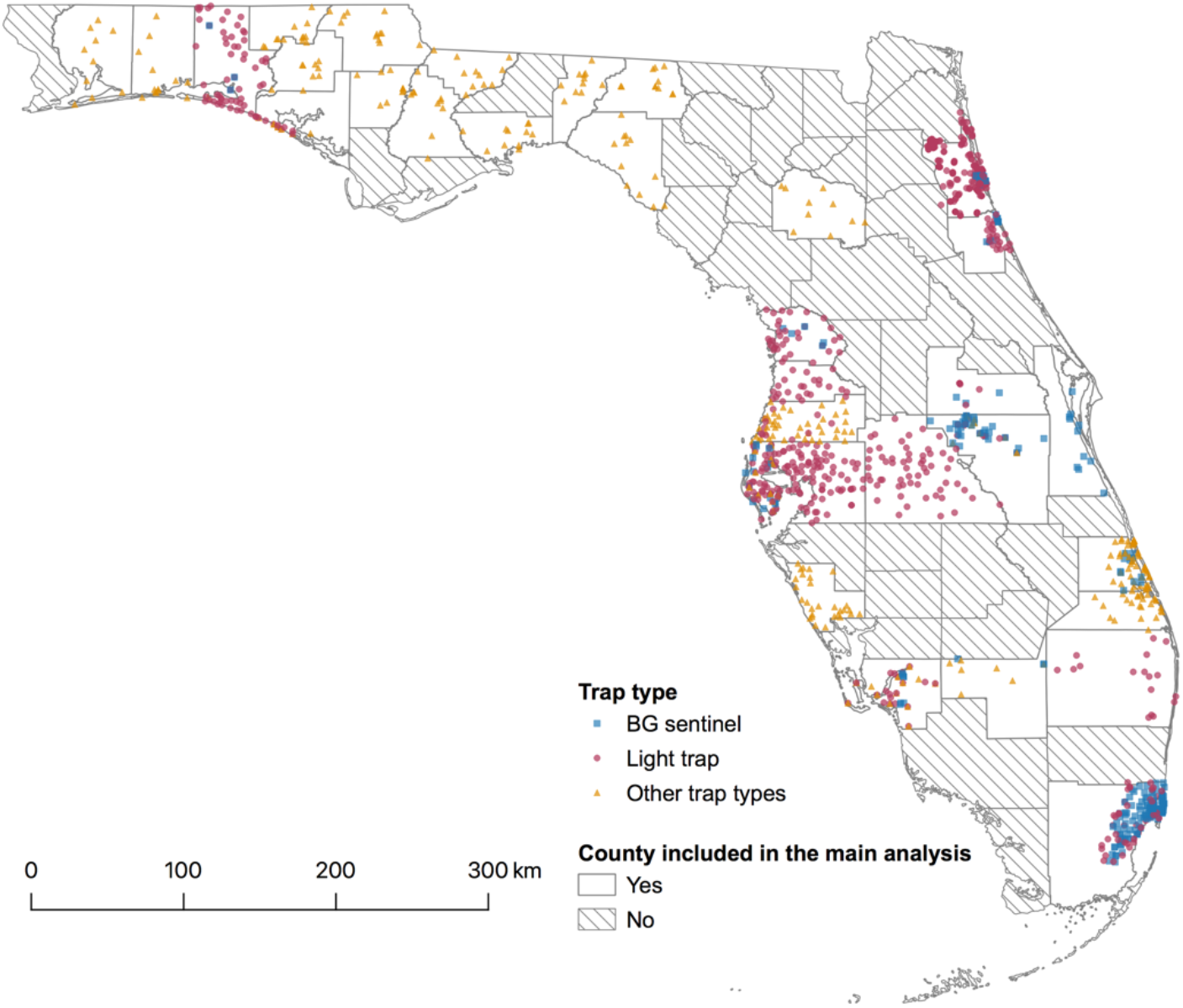
Geographic distribution of mosquito trap types in the longitudinal training dataset.

**Supplementary Figure S9.**
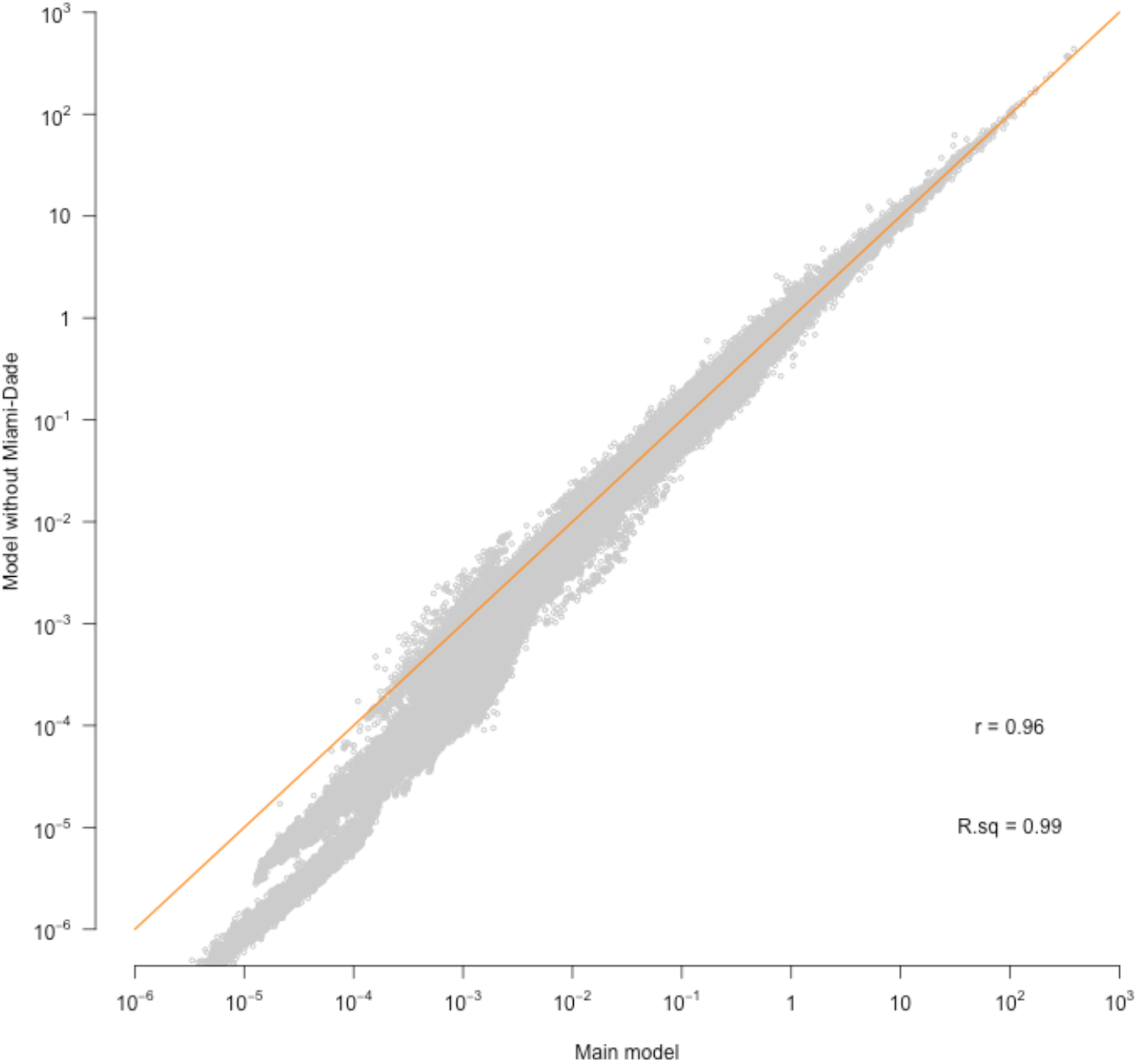
Correlation between predicted trap rate for *Aedes aegypti* using longitudinal data with and without data from Miami-Dade.

**Figure S10.**
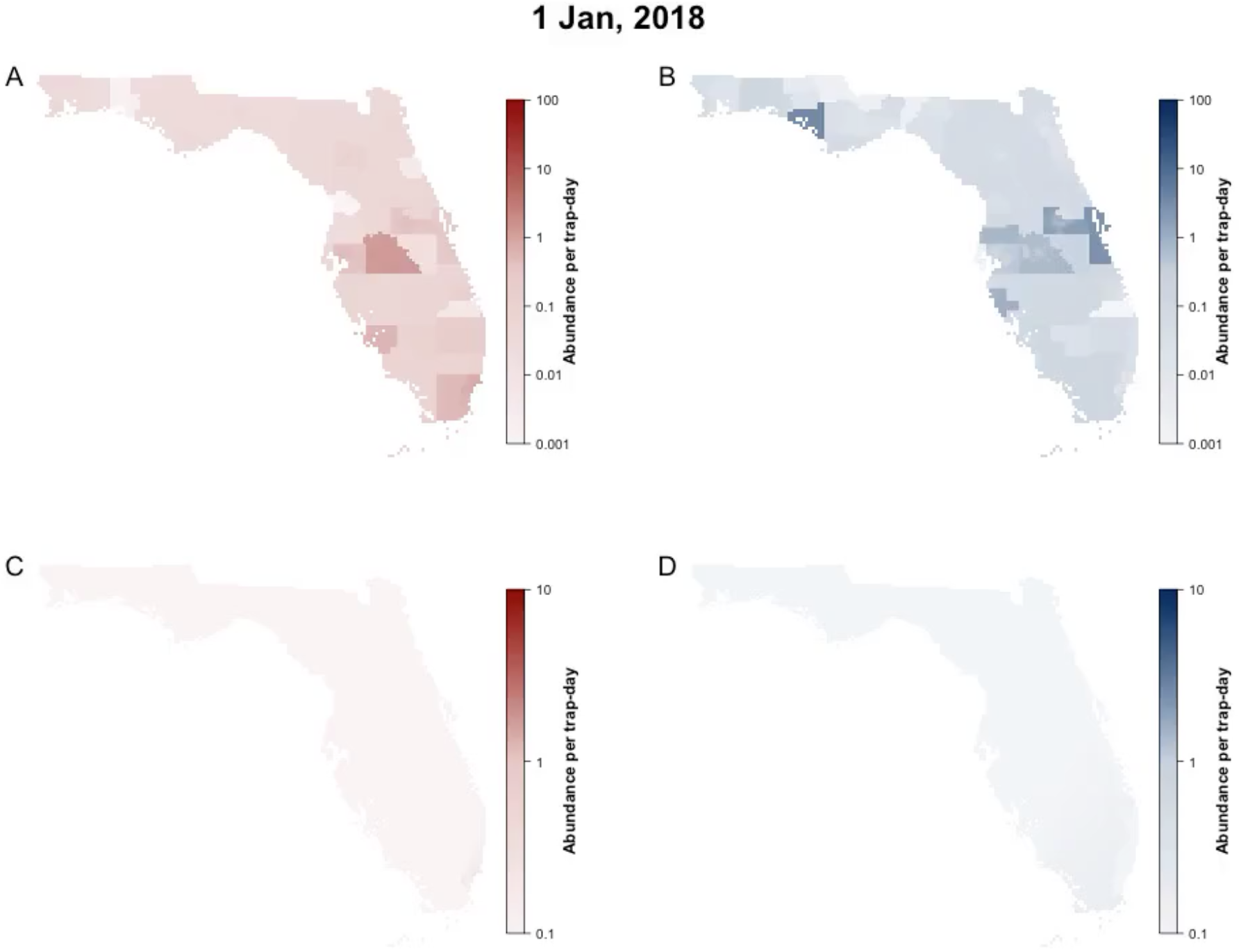
Maps on predicted abundance of *Aedes aegypti* (red) and *Aedes albopictus* (blue) in Florida, 2018. Predictions are derived from “no abundance model”.

**Supplement Table S1.**
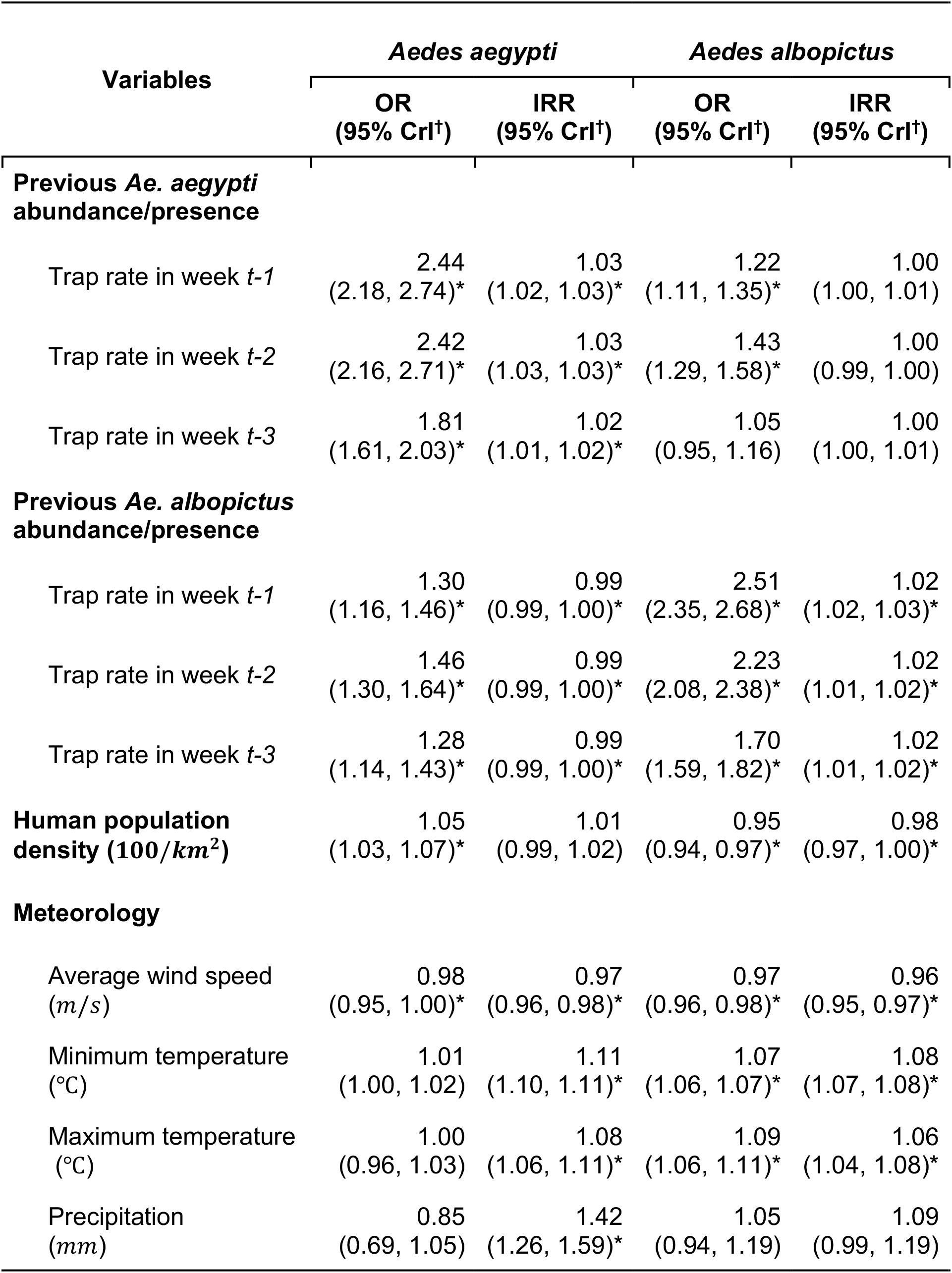
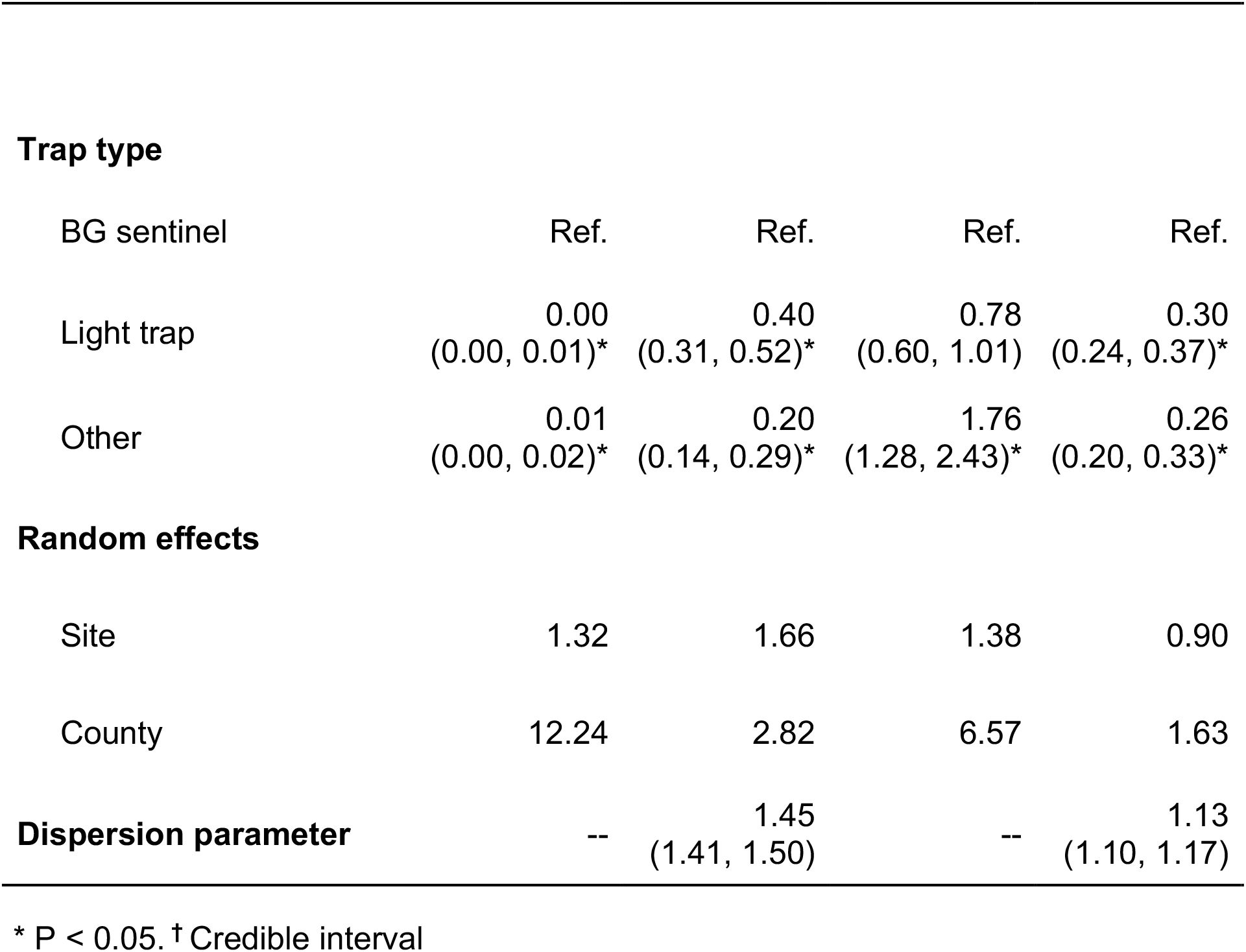
Odds ratio (OR) and incidence rate ratio (IRR) estimate from mixed-effects zero-inflated negative binomial analysis of covariates of *Aedes* trap rates in Florida using data from NOAA, from 2004 to 2018.

| Variables | *Aedes aegypti* |  | *Aedes albopictus* |  |
| --- | --- | --- | --- | --- |
|  | OR<br>(95% CrI<sup>†</sup>) | IRR<br>(95% CrI<sup>†</sup>) | OR<br>(95% CrI<sup>†</sup>) | IRR<br>(95% CrI<sup>†</sup>) |
| **Previous *Ae. aegypti* abundance/presence** |  |  |  |  |
| Trap rate in week *t*-1 | 2.44<br>(2.18, 2.74)* | 1.03<br>(1.02, 1.03)* | 1.22<br>(1.11, 1.35)* | 1.00<br>(1.00, 1.01) |
| Trap rate in week *t*-2 | 2.42<br>(2.16, 2.71)* | 1.03<br>(1.03, 1.03)* | 1.43<br>(1.29, 1.58)* | 1.00<br>(0.99, 1.00) |
| Trap rate in week *t*-3 | 1.81<br>(1.61, 2.03)* | 1.02<br>(1.01, 1.02)* | 1.05<br>(0.95, 1.16) | 1.00<br>(1.00, 1.01) |
| **Previous *Ae. albopictus* abundance/presence** |  |  |  |  |
| Trap rate in week *t*-1 | 1.30<br>(1.16, 1.46)* | 0.99<br>(0.99, 1.00)* | 2.51<br>(2.35, 2.68)* | 1.02<br>(1.02, 1.03)* |
| Trap rate in week *t*-2 | 1.46<br>(1.30, 1.64)* | 0.99<br>(0.99, 1.00)* | 2.23<br>(2.08, 2.38)* | 1.02<br>(1.01, 1.02)* |
| Trap rate in week *t*-3 | 1.28<br>(1.14, 1.43)* | 0.99<br>(0.99, 1.00)* | 1.70<br>(1.59, 1.82)* | 1.02<br>(1.01, 1.02)* |
| **Human population density (100/km<sup>2</sup>)** | 1.05<br>(1.03, 1.07)* | 1.01<br>(0.99, 1.02) | 0.95<br>(0.94, 0.97)* | 0.98<br>(0.97, 1.00)* |
| **Meteorology** |  |  |  |  |
| Average wind speed (m/s) | 0.98<br>(0.95, 1.00)* | 0.97<br>(0.96, 0.98)* | 0.97<br>(0.96, 0.98)* | 0.96<br>(0.95, 0.97)* |
| Minimum temperature (°C) | 1.01<br>(1.00, 1.02) | 1.11<br>(1.10, 1.11)* | 1.07<br>(1.06, 1.07)* | 1.08<br>(1.07, 1.08)* |
| Maximum temperature (°C) | 1.00<br>(0.96, 1.03) | 1.08<br>(1.06, 1.11)* | 1.09<br>(1.06, 1.11)* | 1.06<br>(1.04, 1.08)* |
| Precipitation (mm) | 0.85<br>(0.69, 1.05) | 1.42<br>(1.26, 1.59)* | 1.05<br>(0.94, 1.19) | 1.09<br>(0.99, 1.19) |
| **Trap type** |  |  |  |  |
| BG sentinel | Ref. | Ref. | Ref. | Ref. |
| Light trap | 0.00<br>(0.00, 0.01)* | 0.40<br>(0.31, 0.52)* | 0.78<br>(0.60, 1.01) | 0.30<br>(0.24, 0.37)* |
| Other | 0.01<br>(0.00, 0.02)* | 0.20<br>(0.14, 0.29)* | 1.76<br>(1.28, 2.43)* | 0.26<br>(0.20, 0.33)* |
| **Random effects** |  |  |  |  |
| Site | 1.32 | 1.66 | 1.38 | 0.90 |
| County | 12.24 | 2.82 | 6.57 | 1.63 |
| **Dispersion parameter** | -- | 1.45<br>(1.41, 1.50) | -- | 1.13<br>(1.10, 1.17) |

\* P < 0.05. † Credible interval

**Supplement Table S2.** Odds ratio (OR) and incidence rate ratio (IRR) estimate from mixed-effects zero-inflated negative binomial analysis of covariates of *Aedes aetypti* and *Aedes albopictus* collected from BG traps.

| Variables | *Aedes aegypti* |  | *Aedes albopictus* |  |
| --- | --- | --- | --- | --- |
|  | OR<br>(95% CrI<sup>†</sup>) | IRR<br>(95% CrI<sup>†</sup>) | OR<br>(95% CrI<sup>†</sup>) | IRR<br>(95% CrI<sup>†</sup>) |
| **Previous *Ae. aegypti* abundance/presence** |  |  |  |  |
| Trap rate in week *t*-1 | 5.16<br>(2.45, 10.87)* | 1.04<br>(1.03, 1.04)* | 0.99<br>(0.59, 1.67) | 1.00<br>(0.99, 1.02) |
| Trap rate in week *t*-2 | 6.61<br>(3.48, 12.54)* | 1.02<br>(1.02, 1.03)* | 1.10<br>(0.64, 1.89) | 1.01<br>(0.99, 1.03) |
| Trap rate in week *t*-3 | 8.58<br>(4.13, 17.85)* | 1.02<br>(1.01, 1.02)* | 0.65<br>(0.40, 1.08) | 1.00<br>(0.98, 1.02) |
| **Previous *Ae. albopictus* abundance/presence** |  |  |  |  |
| Trap rate in week *t*-1 | 0.42<br>(0.22, 0.80)* | 1.02<br>(1.00, 1.03)* | 3.27<br>(1.98, 5.41)* | 1.06<br>(1.04, 1.07)* |
| Trap rate in week *t*-2 | 0.96<br>(0.49, 1.90) | 0.99<br>(0.98, 1.01) | 1.12<br>(0.70, 1.79) | 1.06<br>(1.04, 1.07)* |
| Trap rate in week *t*-3 | 0.78<br>(0.41, 1.50) | 0.99<br>(0.98, 1.01) | 3.00<br>(1.89, 4.77)* | 1.04<br>(1.02, 1.05)* |
| **Human population density (100/km<sup>2</sup>)** | 1.12<br>(1.03, 1.23)* | 0.99<br>(0.98, 1.01) | 0.94<br>(0.90, 0.97)* | 0.99<br>(0.96, 1.02) |
| **Meteorology** |  |  |  |  |
| Average wind speed (m/s) | 1.16<br>(0.99, 1.35) | 0.97<br>(0.96, 0.99)* | 1.02<br>(0.92, 1.12) | 0.91<br>(0.87, 0.96)* |
| Minimum temperature (°C) | 1.02<br>(0.93, 1.13) | 1.12<br>(1.11, 1.13)* | 1.02<br>(0.97, 1.07) | 1.10<br>(1.07, 1.12)* |
| Maximum temperature (°C) | 1.25<br>(0.92, 1.70) | 1.07<br>(1.04, 1.10)* | 1.02<br>(0.86, 1.21) | 1.07<br>(0.99, 1.16) |
| Relative humidity (%) | 0.31<br>(0.06, 1.53) | 1.58<br>(1.32, 1.89)* | 0.39<br>(0.18, 0.83)* | 1.27<br>(0.93, 1.73) |
| **Random effects** |  |  |  |  |
| Site | 0.30 | 0.60 | 0.87 | 0.57 |
| County | 2.36 | 0.81 | 4.30 | 1.19 |
| **Dispersion parameter** | -- | 1.35<br>(1.31, 1.40) | -- | 1.10<br>(0.98, 1.23) |

\* P < 0.05. † Credible interval

**Supplementary Table S3.**
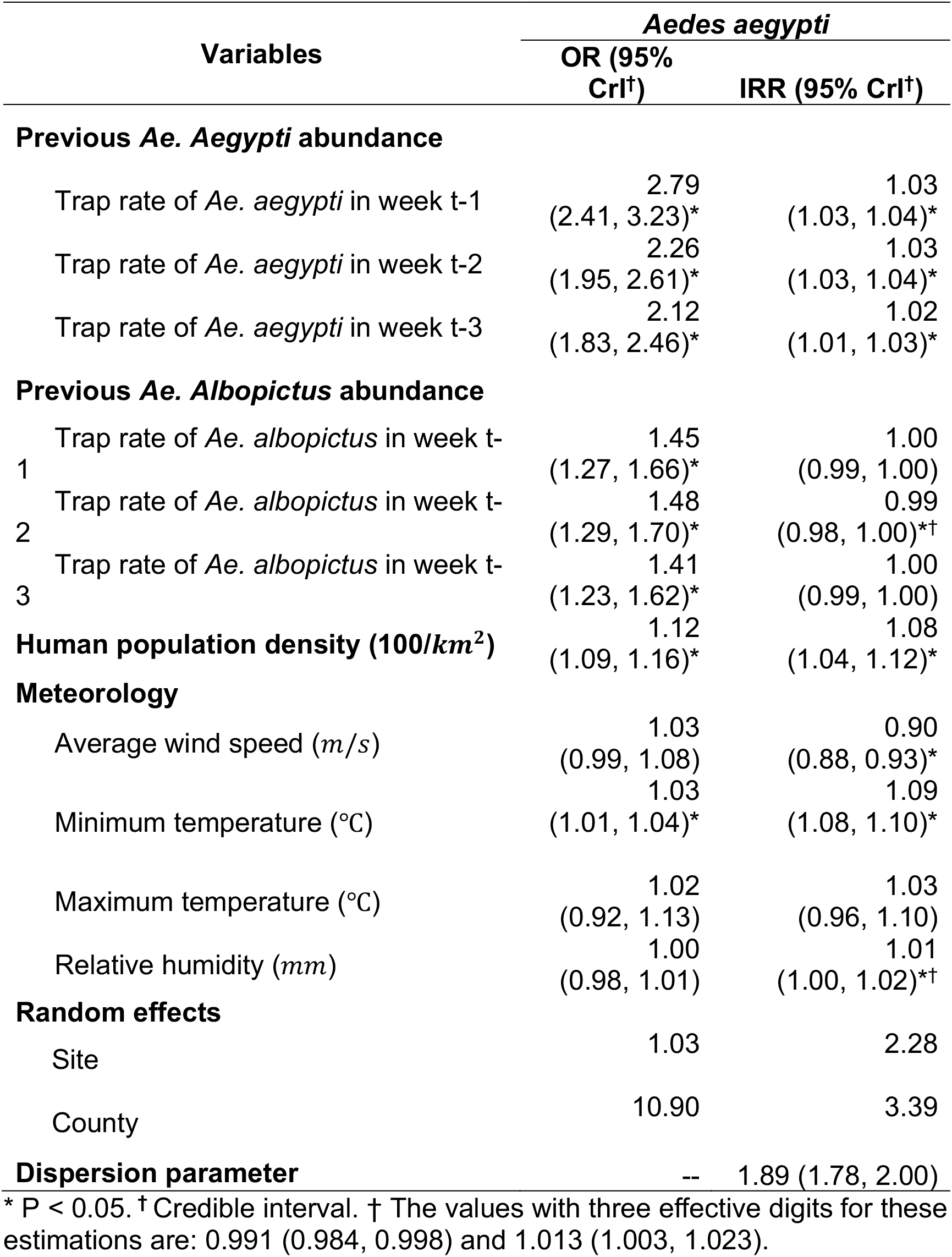
Odds ratio (OR) and incidence rate ratio (IRR) estimate from mixed-effects zero-inflated negative binomial analysis of covariates of *Aedes aegypti* after removing data from Miami-Dade county.

**Supplementary Table S4.** Summary of other trap types included in the longitudinal training dataset.

| Trap types | Number of records |
| --- | --- |
| **BG sentinel trap** | 9,518 (7.2%) |
| **Light trap** | 107,571 (81.4%) |
| CDC light traps | 95,554 (88.8%) |
| New Jersey light traps | 8,451 (7.9%) |
| Non-specific light traps | 3,566 (3.3%) |
| **Other trap types** | 14,999 (11.4%) |
| Mosquito magnet | 3,545 (23.6%) |
| Suction trap | 3,372 (22.5%) |
| Propane | 2,676 (17.8%) |
| ABC | 1,920 (12.8%) |
| Gravid trap | 1,827 (12.2%) |
| Exit | 1,178 (7.9%) |
| Route | 268 (1.8%) |
| Unknown | 139 (0.9%) |
| Fay prince | 64 (0.4%) |
| Wilton trap | 10 (0.1%) |

**Supplementary Table S5.** Surveillance data by county

| County | Full | ZINB | Spatial Training | Temporal Training | No Abundance Testing |
| --- | --- | --- | --- | --- | --- |
| Hillsborough | 24854<br>(14.0%) | 16475<br>(12.5%) | 14861<br>(12.5%) | 16475<br>(12.7%) | 8379<br>(18.4%) |
| Pinellas | 22335<br>(12.6%) | 20058<br>(15.2%) | 18607<br>(15.7%) | 19673<br>(15.2%) | 2277<br>(5.0%) |
| St. Johns | 21872<br>(12.3%) | 17751<br>(13.4%) | 15954<br>(13.4%) | 17751<br>(13.7%) | 4121<br>(9.1%) |
| Polk | 20751<br>(11.7%) | 15543<br>(11.8%) | 12528<br>(10.5%) | 15543<br>(12.0%) | 5208<br>(11.4%) |
| Dade | 18634<br>(10.5%) | 14980<br>(11.3%) | 14129<br>(11.9%) | 14158<br>(10.9%) | 3654<br>(8.0%) |
| Lee | 13812 (7.8%) | 8045 (6.1%) | 6613 (5.6%) | 8045 (6.2%) | 5767 (12.7%) |
| Citrus | 8471 (4.8%) | 6959 (5.3%) | 6695 (5.6%) | 6959 (5.4%) | 1512 (3.3%) |
| Walton | 8380 (4.7%) | 6186 (4.7%) | 5890 (5.0%) | 6106 (4.7%) | 2194 (4.8%) |
| Palm Beach | 7864 (4.4%) | 7008 (5.3%) | 6551 (5.5%) | 6912 (5.3%) | 856 (1.9%) |
| Pasco | 5722 (3.2%) | 3468 (2.6%) | 2922 (2.5%) | 3468 (2.7%) | 2254 (5.0%) |
| Osceola | 4522 (2.5%) | 2203 (1.7%) | 1989 (1.7%) | 2203 (1.7%) | 2319 (5.1%) |
| St. Lucie | 3717 (2.1%) | 2746 (2.1%) | 2527 (2.1%) | 2493 (1.9%) | 971 (2.1%) |
| Flagler | 3715 (2.1%) | 3150 (2.4%) | 2880 (2.4%) | 3118 (2.4%) | 565 (1.2%) |
| Martin | 2660 (1.5%) | 2561 (1.9%) | 2453 (2.1%) | 2350 (1.8%) | 99 (0.2%) |
| Alachua | 2015 (1.1%) | 1538 (1.2%) | 1194 (1.0%) | 1538 (1.2%) | 477 (1.0%) |
| Hernando | 1868 (1.1%) | 1526 (1.2%) | 1299 (1.1%) | 1526 (1.2%) | 342 (0.8%) |
| Hendry | 1052 (0.6%) | 437 (0.3%) | 337 (0.3%) | 437 (0.3%) | 615 (1.4%) |
| Sarasota | 1004 (0.6%) | 268 (0.2%) | 249 (0.2%) | 268 (0.2%) | 736 (1.6%) |
| Bay | 933 (0.5%) | 137 (0.1%) | 105 (0.1%) | 137 (0.1%) | 796 (1.7%) |
| Orange | 724 (0.4%) | 35 (0%) | 35 (0%) | 35 (0%) | 689 (1.5%) |
| Okaloosa | 324 (0.2%) | 216 (0.2%) | 216 (0.2%) | 132 (0.1%) | 108 (0.2%) |
| Santa Rosa | 324 (0.2%) | 168 (0.1%) | 168 (0.1%) | 108 (0.1%) | 156 (0.3%) |
| Brevard | 176 (0.1%) | 30 (0%) | 25 (0%) | 30 (0%) | 146 (0.3%) |
| Holmes | 156 (0.1%) | 84 (0.1%) | 84 (0.1%) | 0 (0%) | 72 (0.2%) |
| Liberty | 150 (0.1%) | 84 (0.1%) | 77 (0.1%) | 0 (0%) | 66 (0.1%) |
| Madison | 149 (0.1%) | 44 (0%) | 40 (0%) | 0 (0%) | 105 (0.2%) |
| Bradford | 137 (0.1%) | 0 (0%) | 0 (0%) | 0 (0%) | 137 (0.3%) |
| Wakulla | 132 (0.1%) | 82 (0.1%) | 82 (0.1%) | 0 (0%) | 50 (0.1%) |
| Indian River | 130 (0.1%) | 0 (0%) | 0 (0%) | 0 (0%) | 130 (0.3%) |
| Washington | 130 (0.1%) | 38 (0%) | 26 (0%) | 0 (0%) | 92 (0.2%) |
| Taylor | 120 (0.1%) | 84 (0.1%) | 84 (0.1%) | 0 (0%) | 36 (0.1%) |
| Gadsden | 108 (0.1%) | 48 (0%) | 44 (0%) | 0 (0%) | 60 (0.1%) |
| Jackson | 108 (0.1%) | 56 (0%) | 53 (0%) | 0 (0%) | 52 (0.1%) |
| Jefferson | 108 (0.1%) | 41 (0%) | 33 (0%) | 0 (0%) | 67 (0.1%) |
| Calhoun | 99 (0.1%) | 39 (0%) | 31 (0%) | 0 (0%) | 60 (0.1%) |
| Collier | 91 (0.1%) | 0 (0%) | 0 (0%) | 0 (0%) | 91 (0.2%) |
| Gulf | 65 (0%) | 0 (0%) | 0 (0%) | 0 (0%) | 65 (0.1%) |
| Charlotte | 57 (0%) | 0 (0%) | 0 (0%) | 0 (0%) | 57 (0.1%) |
| Escambia | 40 (0%) | 0 (0%) | 0 (0%) | 0 (0%) | 40 (0.1%) |
| Okeechobee | 31 (0%) | 0 (0%) | 0 (0%) | 0 (0%) | 31 (0.1%) |
| Union | 15 (0%) | 0 (0%) | 0 (0%) | 0 (0%) | 15 (0%) |
| Leon | 13 (0%) | 0 (0%) | 0 (0%) | 0 (0%) | 13 (0%) |
| Baker | 10 (0%) | 0 (0%) | 0 (0%) | 0 (0%) | 10 (0%) |
| Dixie | 10 (0%) | 0 (0%) | 0 (0%) | 0 (0%) | 10 (0%) |
| Gilchrist | 10 (0%) | 0 (0%) | 0 (0%) | 0 (0%) | 10 (0%) |
| Marion | 10 (0%) | 0 (0%) | 0 (0%) | 0 (0%) | 10 (0%) |
| Suwannee | 10 (0%) | 0 (0%) | 0 (0%) | 0 (0%) | 10 (0%) |
| Nassau | 5 (0%) | 0 (0%) | 0 (0%) | 0 (0%) | 5 (0%) |
| Total | 177623 | 132088 | 118781 | 129465 | 45535 |

